# A method for correcting systematic error in PO_2_ measurement to improve measures of oxygen supply capacity

**DOI:** 10.1101/2024.02.25.581970

**Authors:** A.W. Timpe, B.A Seibel

**Author notes:** For submission to: Comparative Biochemistry and Physiology Part A. **Author Contributions:** Both authors contributed to the conception and writing of this manuscript. AWT developed the methodology and performed laboratory investigations and formal analyses. **Data Availability:** All original data used in this manuscript are available online. Citations are provided for all published data used in this paper.

## Abstract

An organism’s oxygen supply capacity (*α*) can be measured as a ratio of the metabolic rate (MR) and the critical oxygen partial pressure (P_c_). However, this metric is sensitive to errors in the measurement of PO_2_, especially at low PO_2_ where the ratio of instrument error to environmental oxygen is magnified. Consequently, the oxygen supply capacity of animals, particularly those that evolved in low-oxygen environments, may appear unnaturally high and variable. Here, we present a method to correct for instrumental calibration error and use simulated and literature datasets to demonstrate how it can be used *post hoc* to diagnose error in, correct the magnitude of, and reduce the variability in repeat measures of the oxygen supply capacity.

## Introduction

Low environmental oxygen is a common challenge for aquatic organisms. In large regions of the world’s oceans, including oxygen minimum zones (Fuenzalida et al., 2009; Wyrtki, 1962), continental margins (Helly and Levin, 2004), at the outflow of eutrophic river systems (i.e., “dead zones;” Rabalais et al. 2002), and in estuaries, tidepools and stagnant freshwater systems (Seibel, 2024a, 2024b), animals routinely experience oxygen partial pressures (PO_2_) well below air saturation. Although there is abundant life in low-oxygen areas (Childress and Seibel, 1998; Levin, 2002), survival there requires specific adaptations for oxygen uptake and transport. Thus, oxygen variability in the ocean is an important parameter that may limit growth, reproduction, activity and biogeography (Breitburg et al., 2018; Richards, 2011; Seibel, 2024a). Moreover, due to anthropogenic influences, oxygen is declining globally (Breitburg et al., 2018; Jenny et al., 2016; Keeling et al., 2010; Oschlies, 2021).

A common method to quantify an animal’s metabolic response to changes in or different levels of environmental oxygen is respirometry, which measures an organism’s oxygen uptake rate. During a respirometry trial, the rate of oxygen consumption is calculated in discreet bins, defined by time or PO_2_ (Steffensen, 1989; Svendsen et al., 2016), and is a commonly-used indirect measure of whole-animal aerobic metabolic rate (Killen et al., 2021; Nelson, 2016; Steffensen, 1989). The response of metabolic rate to changes in available oxygen inform physiological metrics such as the critical oxygen partial pressure (P_c_; Farrell and Richards, 2009) and oxygen supply capacity (α; Seibel and Deutsch, 2020).

These indices can supplement measures of standard (SMR; Chabot et al., 2016; Claireaux and Chabot, 2016) and maximum metabolic rate (MMR; Norin and Clark, 2016) to address ecological and biogeographical questions.

Though there has been lively recent debate of the definition and utility of these metrics (Farrell et al., 2021; Seibel et al., 2021b, 2021a; Seibel and Deutsch, 2020; Wood, 2018), both α (or P_c_) and MR must be determined with high precision and accuracy if they are to be of any utility. Killen et al. (2021) noted that inconsistent methods and insufficient method reporting imposes challenges when interpreting respirometry data and evaluating its validity. Oxygen probe calibration details were particularly lacking and, while the manufacturers of the wide range of oxygen sensors and probes report high accuracy and precision when properly calibrated (Appendix 1), Helm et al. (2018) observed that standard laboratory practices during sensor calibration can easily introduce small amounts of systematic error in PO_2_ measurement. This type of error can be difficult to diagnose but must be present if negative PO_2_ values are recorded during an experiment or if a positive oxygen consumption rate is recorded at anoxia.

Seibel et al. (2021) recently developed a novel method to directly determine the oxygen supply capacity, defined as the ratio of MR to the P_c_ for that rate (*α* = MR/P_c_). Interestingly, because this method relies on a ratio, it is sensitive to error in the denominator that is introduced during calibration. This is particularly true at low PO_2_ where the ratio of error to environmental oxygen is amplified. Consequently, where PO_2_ measurement error exists, *α* may appear unnaturally high and extremely variable, especially for species with low P_c_. In such instances, the calculated P_c_ may not be representative of the respirometry data or make sense in an ecological context. However, this sensitivity to calibration error provides an opportunity to identify, quantify, and therefore, correct it. Here we present a method for correcting PO_2_ measurement error in respirometry data to improve measurements of oxygen supply and critical oxygen levels.

### Theory and Calculation

The ratio of MR to the PO_2_ for a given measure period is termed the instantaneous oxygen supply, *α*_0_, which typically increases as PO_2_ declines given a relatively constant, “regulated”, oxygen consumption rate (Seibel et al., 2021b). This occurs because the oxygen uptake and transport rates increase as PO_2_ declines to meet a given oxygen demand. In other words, per unit of available oxygen, more oxygen is taken up as PO_2_ declines because ventilation and heart rate increase. The oxygen supply capacity, *α*, is the maximum *α*_0_value and is reached at the P_c_ for the coincident metabolic rate. This method is not affected by changes in MR caused by diel cycles or spontaneous activity because *α* is independent of the rate at which it is determined.

To visualize the effect of systematic PO_2_ measurement error (S) on *α*_0_, we created a simulated dataset in which a known amount of error was added uniformly to all PO_2_values. The instantaneous oxygen supply with error 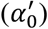 for each (PO_2_, MR) pair was calculated as 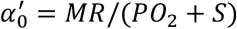. As PO_2_ approaches zero, the influence of such error on the calculated *α*_0_ increases dramatically. If measured PO_2_ is an underestimate of reality (e.g., negative PO_2_ values are measured due to a high zero calibration), 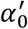 diverges toward infinity as PO_2_ approaches zero (Fig. 1). The method we present here exploits that divergence to identify calibration error. Correction of that error relies on the assumption that, if the effect of error were removed, there would be two bins where *α*_0_ = *α*. Even if *α*_0_ falls immediately upon reaching P_c_, this assumption can be met by adjusting the MR averaging period (bin size) and imposes no unique requirements on the respirometry methodology.

**Figure 1.**
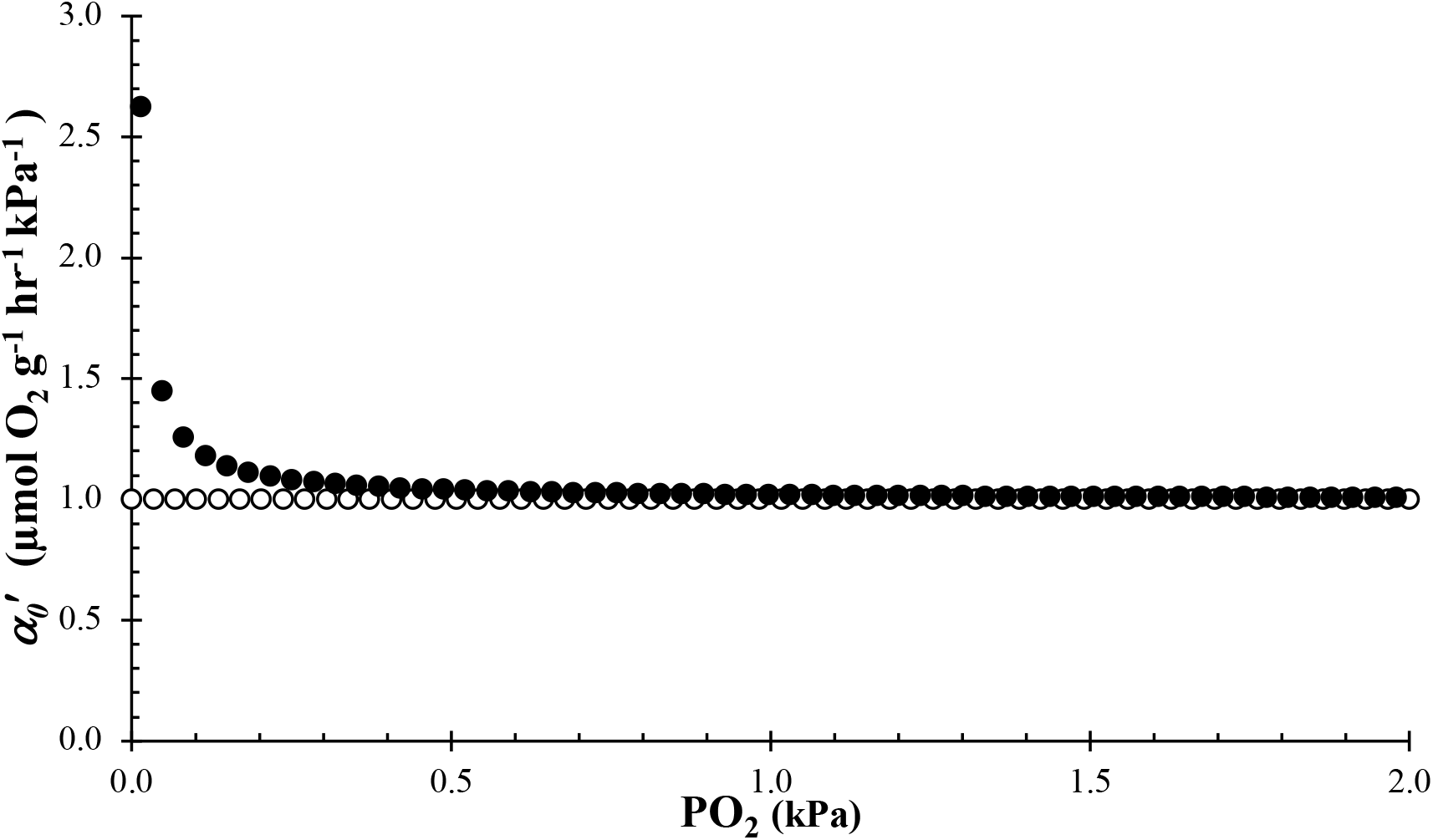
Example maximum metabolic rate dataset (MMR; *α*_*0*_ *= α*) showing the divergence of *α*_0_ near anoxia for data with -0.1% (−0.021 kPa) systematic error (*S*) in PO_2_ measurements (black circles). Data without introduced systematic error (white circles) are provided for reference. The input *α* value for the dataset was 1.0.

To correct for systematic error (*S*) in PO_2_ measurement, a respirometry trial must include two or more measurement periods during which the animal’s oxygen supply (*α*_0_) is operating at maximum capacity (*α*). Error is estimated by identifying discontinuities in the second derivative of *α* with respect to a PO_2_ correction value (*C*, defined below). At such discontinuities, two (PO_2_, MR) points provide equivalent estimations of *α*. Here we demonstrate, using simulated and literature datasets, that correcting respirometry data in the manner described successfully eliminates the effect of PO_2_ calibration error on the calculated value of *α*. The following calculations can be performed using an R or Excel template available in the online supplementary materials for this manuscript. A summary reference for the notation used in this manuscript is provided in Table 1.

**Table 1.** List of Metrics and Variables.

| Symbol | Name | Units |
| --- | --- | --- |
| $\alpha$ | Physiological oxygen supply capacity | $\mu\text{mol O}_2 \text{ g}^{-1} \text{ hr}^{-1} \text{ kPa}^{-1}$ |
| $\alpha_0$ | Physiological oxygen supply | $\mu\text{mol O}_2 \text{ g}^{-1} \text{ hr}^{-1} \text{ kPa}^{-1}$ |
| $\alpha'_0$ | Measured physiological oxygen supply | $\mu\text{mol O}_2 \text{ g}^{-1} \text{ hr}^{-1} \text{ kPa}^{-1}$ |
| $\alpha''_0$ | Corrected physiological oxygen supply | $\mu\text{mol O}_2 \text{ g}^{-1} \text{ hr}^{-1} \text{ kPa}^{-1}$ |
| $C$ | Correction value of $\text{PO}_2''$ | kPa |
| $C'$ | Potential correction value for $\text{PO}_2'$ | kPa |
| $C'_i$ | The $i^{\text{th}}$ interval between $\pm C'$ of width $\Delta C_i$ | kPa |
| $\Delta C_i$ | Interval width between potential correction values | kPa |
| $f(C'_i)$ | Maximum physiological oxygen supply at some $C'$ | $\mu\text{mol O}_2 \text{ g}^{-1} \text{ hr}^{-1} \text{ kPa}^{-1}$ |
| MMR | Maximum Metabolic Rate (maximal activity) | $\mu\text{mol O}_2 \text{ g}^{-1} \text{ hr}^{-1}$ |
| MR | Metabolic rate | $\mu\text{mol O}_2 \text{ g}^{-1} \text{ hr}^{-1}$ |
| $P_c$ | Critical oxygen partial pressure | kPa |
| $\text{PO}_2$ | Environmental oxygen partial pressure | kPa |
| $\text{PO}'_2$ | Measured environmental oxygen partial pressure | kPa |
| $\text{PO}''_2$ | Corrected environmental oxygen partial pressure | kPa |
| $S$ | Systematic or calibration error in $\text{PO}_2$ measurement | kPa |
| SMR | Standard Metabolic Rate (resting, fasted animal) | $\mu\text{mol O}_2 \text{ g}^{-1} \text{ hr}^{-1}$ |
| $t$ | Time | Hours |

#### Part I: Two Datapoints

Assume a PO_2_ measurement has some amount of systematic error (*S*) caused by inaccurate sensor calibration. Define the measured oxygen partial pressure, PO_2_′, as:

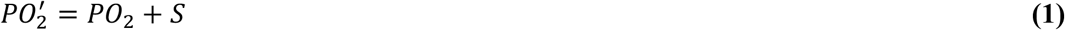

where PO_2_ is the actual oxygen partial pressure without error. Here we assume error is invariant of PO_2_. For a discussion of alternative cases, see Appendix 4. An animal’s aerobic metabolic rate (MR) is then calculated as the negative change in measured oxygen partial pressure over the change in time, *t*, and normalized for chamber size, animal mass, and the number of moles/L at maximum saturation to give final units of μmol/g/hr (Steffensen, 1989). This normalization is omitted below for simplicity. The change in oxygen is negative by convention to provide a positive consumption rate, interpreted as the “amount of oxygen consumed per gram of animal per unit time.”

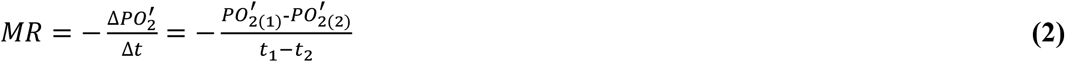

Using (1), (2) can be re-written and simplified:

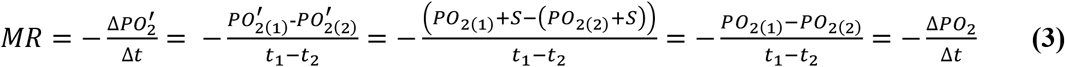

Thus, as evident in (3), systematic error does not bias estimates of metabolic rate. Seibel et al. (2021b) calculate the physiological oxygen supply (*α*_0_) for a given aerobic metabolic rate and level of environmental PO_2_ as:

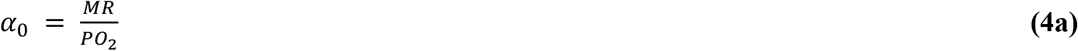

At the critical PO_2_ for a given MR, physiological oxygen supply reaches maximum capacity (*α*), MR is the maximum achievable rate at that PO_2_, and *α*_0_ = *α* (Eq. 4b).

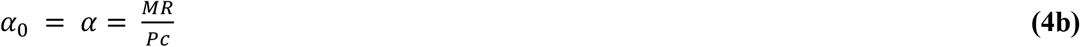

When an organism is operating at its physiological oxygen supply capacity, metabolic rate is dependent upon and linearly proportional to available environmental oxygen (Seibel et al., 2021b, 2021a). If there is calibration error in the observed value of PO_2_, we define the measured physiological oxygen supply, 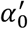, by incorporating (1) into (4a):

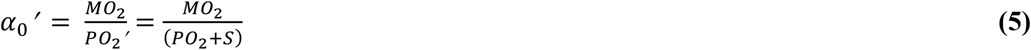

Next, we define a correction factor, *C*, where

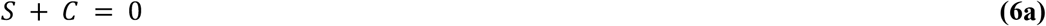

If we add the correction factor *C* to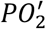, we can define the corrected environmental oxygen partial pressure 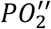 as

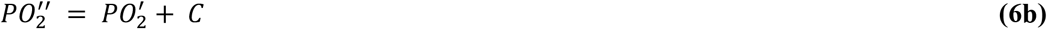

And, using (1), (6b) can be expanded to show:

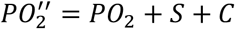

Solving for PO_2_ gives:

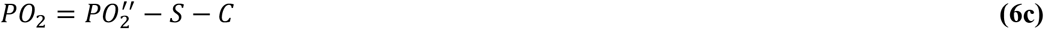

Thus, by (1) and (6a), it can be shown using (6c) that:

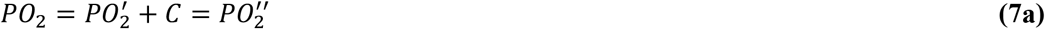

In analogous fashion to our definition of 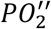, we define the corrected physiological oxygen supply, 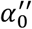, by substituting (7a) into (4a):

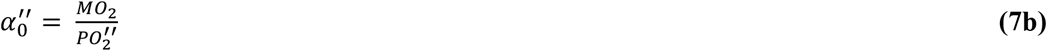

If the conditions of (4b) are satisfied, then by equations (7b), (6b), and (1),

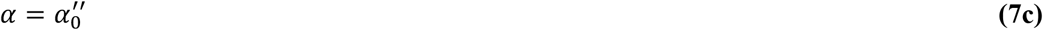

Consider two points, 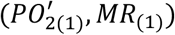 and 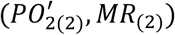where *α* _0 (1)_ = *α* _0 (2)_ = *α*. By (4a) and (4b):

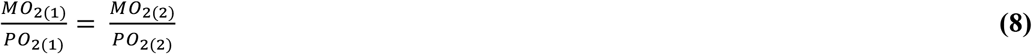

Equivalently, using (6a),

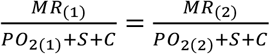

And by (1),

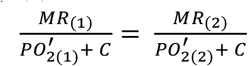

Rearranging allows us to calculate *C* exactly if *MR*_(1)_ ≠ *MR*_(2)_ and 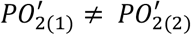:

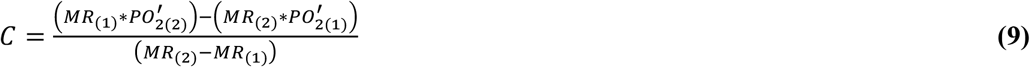

Because *MR*_(*n*)_ and 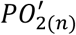 are measured experimentally, we can determine the amount of error and appropriate correction value using (9).

#### Part II: Correcting error in datasets with more than two points

In Part I, we show that systematic error (*S*) in the measurement of oxygen partial pressure (PO_2_) does not affect the calculation of metabolic rate (MR). We also show how the magnitude of such error can be determined if the actual oxygen supply (*α*_0(*n*)_) of two measured points, 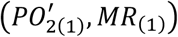 and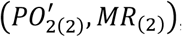, are, without error, at capacity and therefore equal. While this correction is straightforward with two points, the challenge in larger datasets is determining which points should be adjusted such that, after their correction, 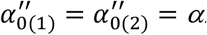. In other words, we must identify observations that would give equivalent estimates of *α* if calibration error were removed. This is particularly important during trials where, for some duration, the animal is operating below capacity (i.e., PO_2_> P_c_). The two points that should be used in eq. 9 are identified by approximating the second derivative of corrected physiological oxygen supply 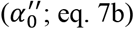 with respect to potential correction value (*C′*). Discontinuities in the second derivative indicate correction values that give equivalent estimates of *α* for two corrected observations.

To correct for error in larger datasets, first create an array of potential correction values with a number, *i*, of discrete, equal intervals of width Δ*C′* between some maximum and minimum range of potential correction values (*±C′*). For example, one might choose an initial range that encompasses -10% to 10% of the saturation value (±2.1kPa). In every test case, *i* = 200 intervals between these boundary values of *C′* provided sufficient resolution to identify the discontinuities used to determine *C*. Furthermore, it is possible to achieve higher resolution by narrowing the boundary conditions or increasing the number of intervals if the initial conditions are insufficient. Next, calculate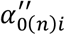, which describes the corrected oxygen supply for the *n*^th^ paired (PO_2_, MR) observation in a respirometry dataset and where 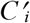 is the value of *C* at the *i*^th^ interval between *±C′* of width Δ*C′* :

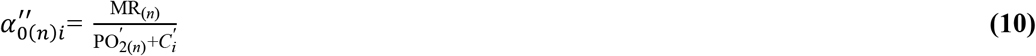

We then plot 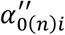 as a function of *C′*. An idealized dataset where the animal always operates at capacity (i.e., is always at MMR) provides the simplest example of the 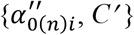 relationship (Fig. 2). Since *α*_*0*_ is maximized at *α* (Seibel et al., 2021b), only the maximum 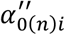 for any point(s) in a respirometry dataset 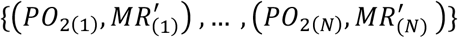 for each 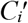 is relevant as only the highest 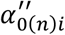 could potentially be at capacity. Moreover, when the two highest 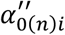 are equal, the conditions of eq. 4b are satisfied and, at that value of *C′*, by eq. 6a, 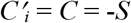. Therefore, to determine where intersections between maximized 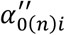 occur, we define a new function 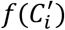 using eq. 10, that describes the maximum 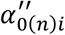 as a function of *C′* : *α*_0_

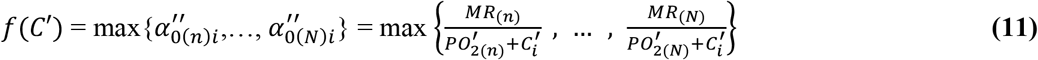

**Figure 2.**
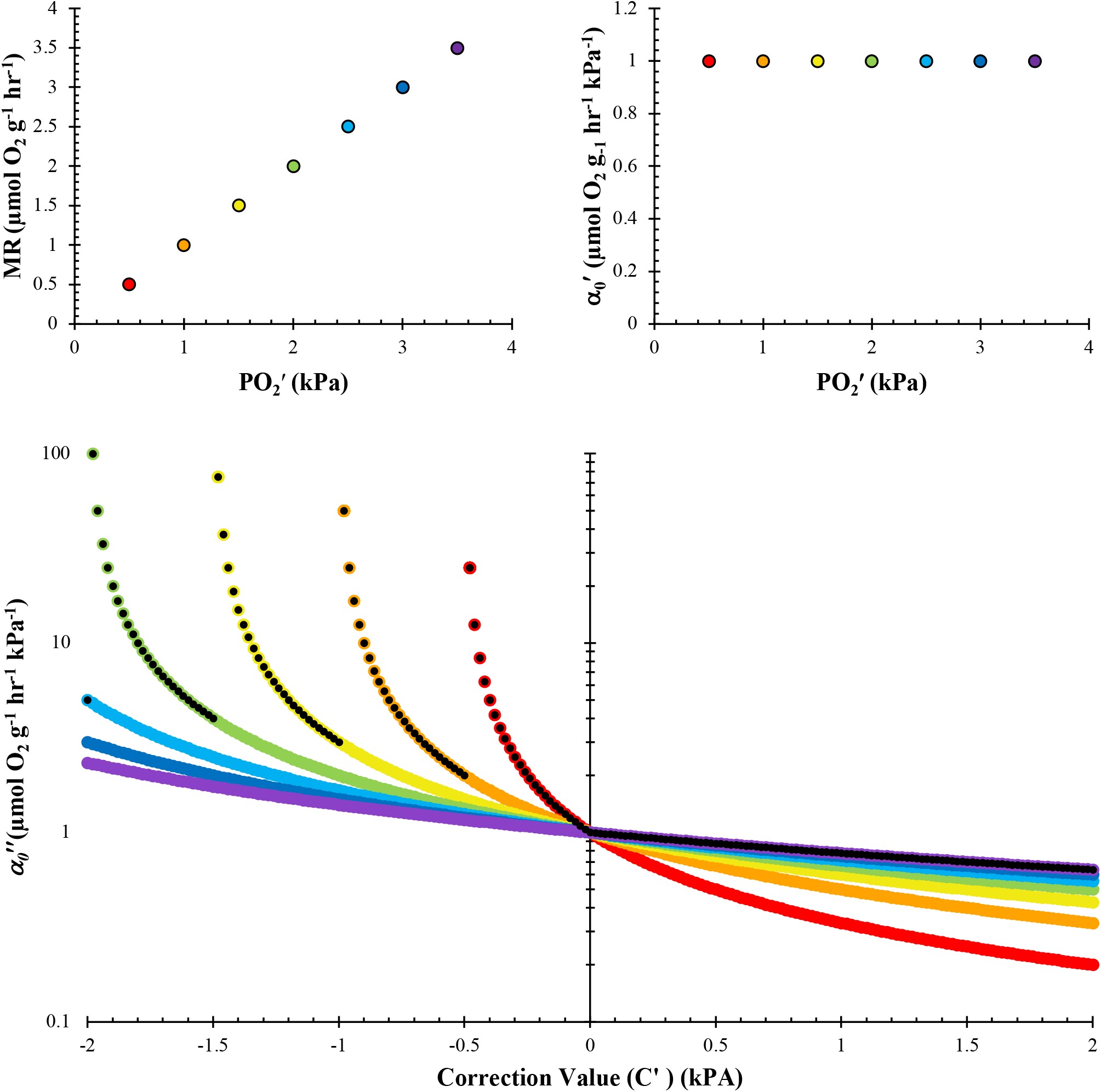
**A)** Example maximum metabolic rate (MMR) dataset 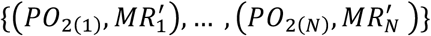 where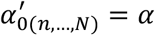. The selected *α* for this dataset = 1.0 and there is no systematic error 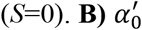 as a function of 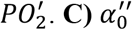 as a function of *C′* at intervals *C′* = 0.02 kPa for all points in the dataset in panel **A** and are color-coded to match. The superimposed black dots identify the points used to define the piecewise function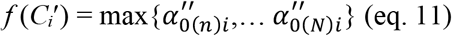. A corrected supply measurement 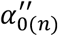 becomes negative when *C′* > PO_2(n)_ and is not displayed on a log-scale. Note that, in this dataset where *S =* 0, the only solution to 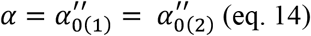 occurs at *C′* = *C* = -*S* = 0 and therefore indicates no correction is needed. An example with error is provided in Appendix 2.

Because there are discontinuities in *f* (*C′*), the function is not continuous for all *C′*. Additionally, at the *C′* where the datapoints that provides the solution to eq. 11 change, *f* (*C′*) may be continuous but not differentiable. Consequently, there are discontinuities in its derivative that identify those asymptotes and intersections. For continuous segments of the function *f* (*C′*) from 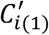 to 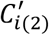 where (PO_2(n)_, MO_2(n)_) provides the solution to eq. 11, *f* (*C′*) is differentiable within that region and

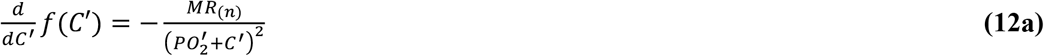

All solutions to eq. 12a must be ≤0 since MR must be ≥0 in heterotrophic organisms and since 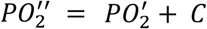 (eq. 6b) must be positive if 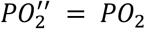 and 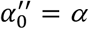. Differentiating again with respect to *C′* in that continuous, bounded region of *C′* gives:

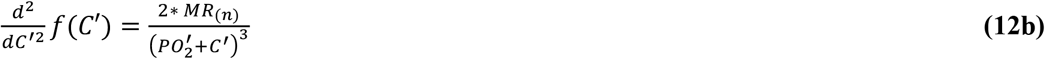

Since the constraints on MR and PO_2_ enumerated above still apply, all valid second derivatives of *f* (*C′*) must be positive. While it would be sufficient to approximate the first derivative to identify discontinuities in *f* (*C′*), they are more obvious using the second derivative as it magnifies the y-axis difference between the continuous, bounded regions of *f′*(*C′*). Additionally, as discontinuities in the second derivative occur at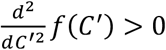, they can be identified in practice as a shift in 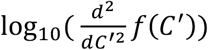 of typically ∼0.3 log units or greater (Fig. 3). Because this function is not differentiable everywhere, it is possible to approximate the second derivative of *f* (*C′*) (eq. 11) at the *i*^th^ interval between ±*C’* of width *C′* by using the second-order central difference quotient:

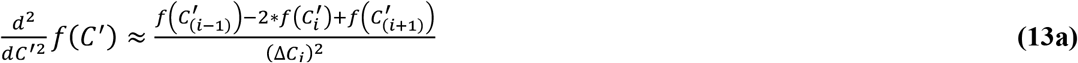

**Figure 3.**
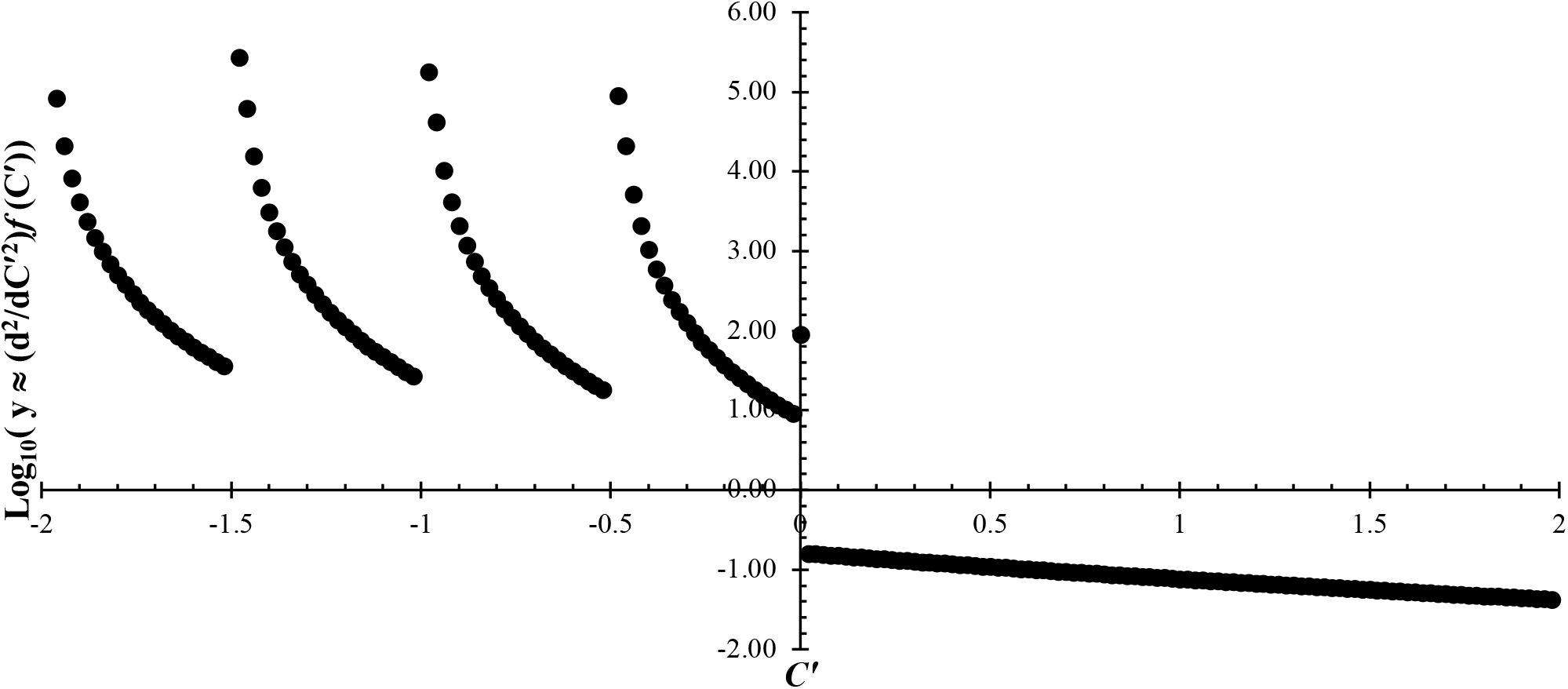
log_10_-transformed second-order difference quotient of *f* (*C′*) with respect to *C′* for the MMR dataset presented in Fig. 2. Discontinuities where potential *C′* = *C* = −*S* are identified by a 0.3 log-unit or greater difference between 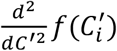 and 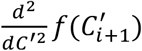 or 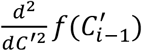, and where 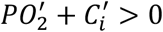 for all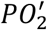. For this dataset where *S* = 0, the only appropriate discontinuity occurs at *C′* = 0 and indicates no correction is needed. Discontinuities at negative *C′* would result in some 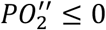 and are therefore invalid.

Equivalently, using eq. 11,

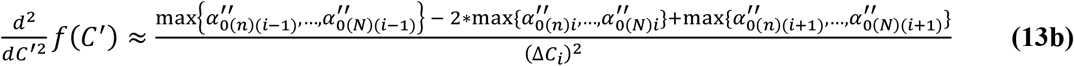

The correction value *C* for the two points, 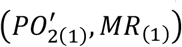 and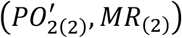, which are the solutions of eq. 13b that bound the discontinuity, is then precisely calculated using eq. 9 and, by eq. 7c and assuming eq. 4b is true for those two points,

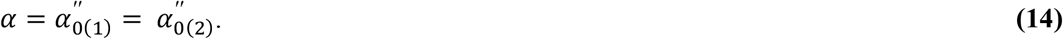

In practice there will likely be more than one discontinuity in *f′′*(*C′*), especially if *C′* is large or there are many observations. Methods for selecting the most appropriate correction value are discussed below.

### Methods: Testing the Correction Method

#### Respirometry

When an animal is operating at its oxygen supply capacity, the amount of oxygen it uses declines in proportion to the amount available (MR = *α*\*P_c_). This can be simulated as the rate of oxygen displacement due to bubbling with nitrogen gas. The rate of displacement varies in proportion to the flow of nitrogen gas. To test the effects of probe calibration error on respirometry data empirically, we purposefully calibrated probes incorrectly, corrected the data, and compared the results to probes calibrated precisely to zero. Precise zero calibrations were performed by immersing oxygen dipping probes (*n* = 3-4; Robust Oxygen Probe OXROB10 or Trace Range Robust Oxygen Probe TROXROB10, PyroScience GMBH, Aachen, Germany) for 30 minutes in a 500mL flask containing a 1%w/v solution of Na_2_SO_3_ that was bubbled continuously with 300cc/min of Nitrogen gas (Airgas USA, Clearwater, FL, USA) through a glass diffuser and regulated by a mass flow controller (Sierra Instruments, Monterey, CA, USA). The flask was covered with Parafilm™ (Amcor, Zurich, Switzerland), immersed in a LaudaE100 temperature-controlled water bath (Lauda-Brinkman LP, Marlton, NJ, USA), and stirred continuously.

Imprecise probe calibrations were achieved by placing precisely-calibrated probes in a solution bubbled with nitrogen until it reached the goal oxygen partial pressure (∼0.05-0.2 kPa) at which point a new, inaccurate low calibration was performed. These inaccurately-calibrated probes were replaced into the original anoxic solution and their error (negative oxygen partial pressure) was recorded for 10 minutes. All probes then recorded the decrease in oxygen of seawater in 500mL flask bubbled with N_2_. Nitrogen bubbling rates across replicate trials ranged from 30 to 500cc/min and was regulated by a mass flow controller (Sierra Instruments, Monterey, CA, USA), but rates were kept constant during each trial to mimic an animal operating at MMR.

Oxygen traces from these experiments were analyzed as if they were respirometry data from an animal, using R (R Core Team, 2024) and the package respirometry v1.2.0 (Birk, 2020). The O_2_ displacement rate, our MR analog, was calculated using sequential bins (Prinzing et al., 2021) and a bin duration that gave approximately the same number of observations (about 40-60) per trial. Each oxygen trace was assessed using the correction method described here. Because N_2_ flow rates differed between trials, data were normalized per trial by dividing the observed “oxygen supply capacity” for each probe with error by the mean “supply capacity” of reference probes without calibration error. Thus, positive values indicate the apparent *α* from probes with calibration error is higher than accurately-calibrated probes and a value of 1 post-correction indicates the method accounted for the effect of calibration error on α. Representative curves from this experiment are provided in Appendix 3.

#### Application to literature datasets

To test the applicability of this method, we evaluated several respirometry datasets. These include “representative curves” for a diverse range of taxa that were extracted from published manuscript figures using WebPlotDigitizer 4.6 (Rohatgi, 2022; https://automeris.io/WebPlotDigitizer/index.html; last accessed Feb 25, 2024) and published datasets for the Pacific oxygen minimum zone copepod *Megacalanus* spp. (Wishner et al., 2018). We also conducted experiments using bubbled N_2_ to simulate an organism operating at maximum capacity (see above). Statistical analyses were conducted using the Paleontological Statistics (PAST) software package v. 4.15 (Hamner et al., 2001). Nonparametric tests were employed when assumptions of normality or homogeneity of variance were rejected.

### Results and Application

#### Appropriate use of the correction method

Calculating the oxygen supply capacity (*α* = MMR/P_crit_; Seibel and Deutsch, 2020) allows for the determination of the critical oxygen partial pressure (P_c_) for any metabolic rate. However, small negative calibration errors in oxygen probes (i.e., less oxygen is measured than is really in solution) can cause the estimate of *α* to be exaggerated and variable among replicate respirometry trials. Therefore, these calibration errors must be recognized and corrected for when they occur.

The effects of negative PO_2_ error on *α* may manifest in the (PO_2_′, *α*) profile three different ways. These profile morphologies illustrate the criteria for appropriate use of the correction method presented here. First, negative PO_2_ recorded during a respirometry trial indicates calibration error and consequently such trials require correction. Such trials can produce extremely variable and high *α*_0_ values that result in ecologically irrelevant estimations of *α* and, thus, P_c_ (Fig. 4). If oxygen is not consumed until *S* > PO_2_′, negative PO_2_ measurements will not be observed despite the presence of calibration error. If *S* is small relative to P_c_, *α*_0_ may diverge at PO_2_ < P_c_. This appears graphically as a plateau in *α*_0_ followed by an increase at low PO_2_ (Fig. 5). We can offer no satisfactory physiological explanations for this type of oxygen supply response and, therefore, recommend employing the correction method for profiles of this type.

**Figure 4.**
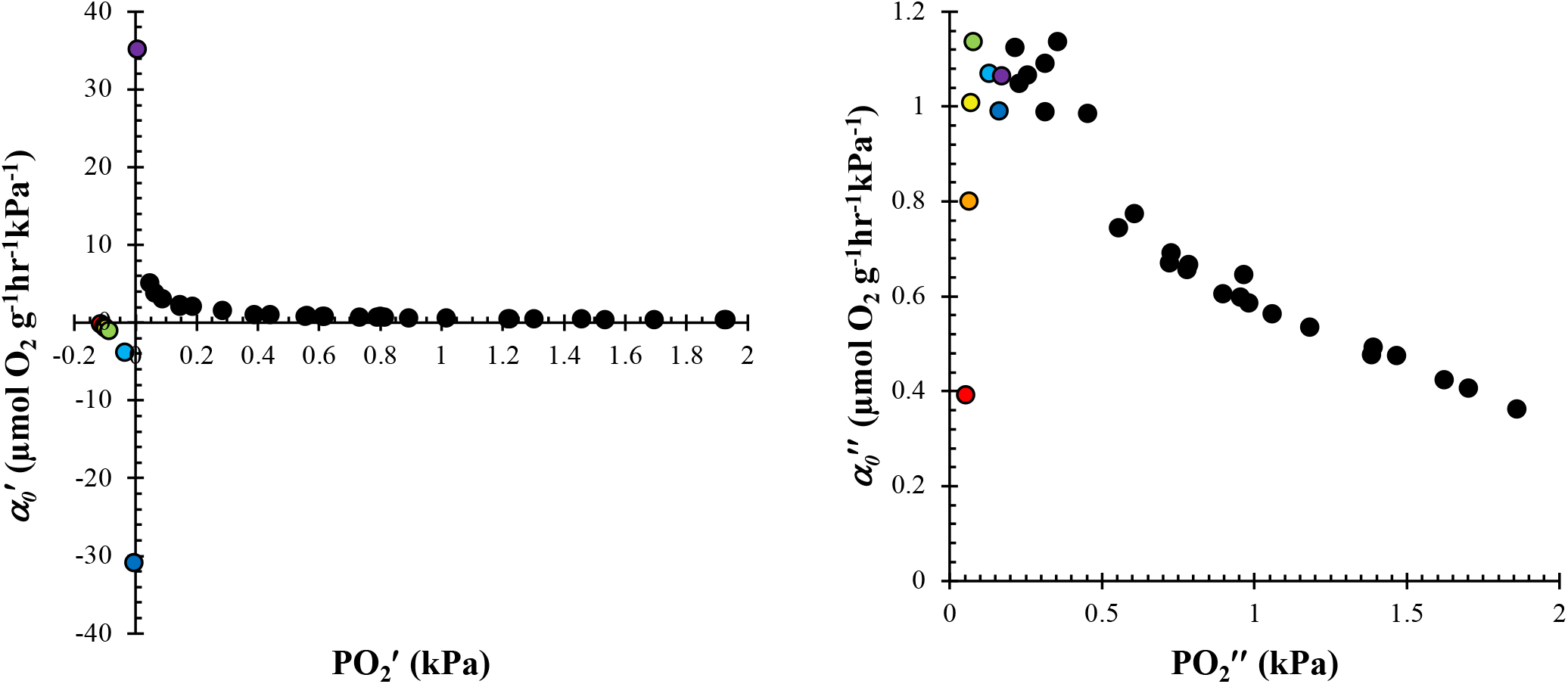
An example PO_2_, *α* curve for the OMZ copepod genus *Megacalanus* (Wishner et al., 2018) in which negative PO_2_ were recorded. **A)** Without correction, negative PO_2_ generates a highly variable 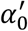 with no discernable pattern and results in an extremely low P_c_ for its routine metabolic rate (P_c_ = 0.024 kPa. **B)** Post-correction (*C* = 0.1663 kPa), all PO_2_ are positive, 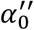 has an interpretable relationship with respect to PO_2_, and P_c_is ecologically relevant at 0.73 kPa. Datapoints in panels A, B are color-coded to match.

**Figure 5.**
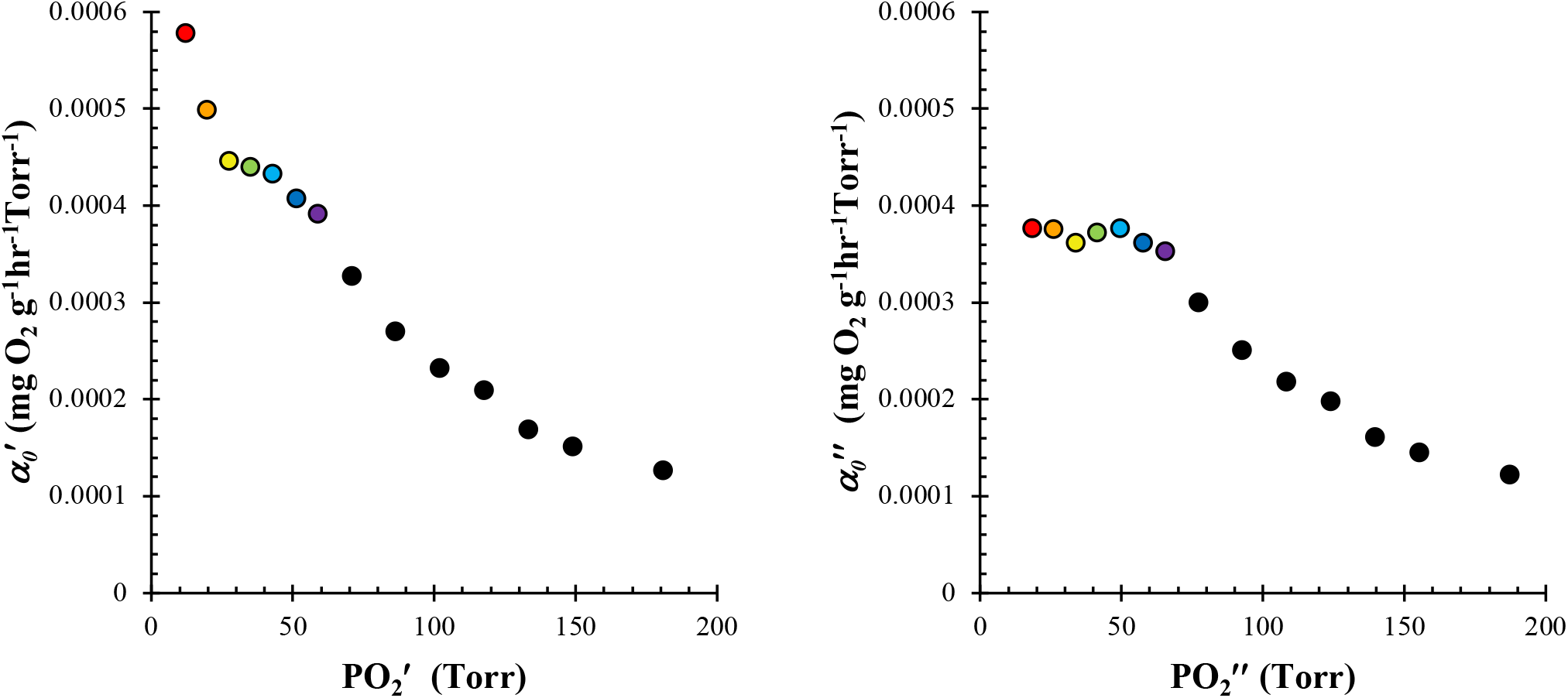
An example PO_2_, *α* curve for the southern rock lobster *Jasus edwardsii* (Crear and Forteath, 2000) illustrating **A)** a plateau in 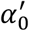 at intermediate PO_2_ followed by an increase at the lowest-recorded values, and **B)** how correcting the dataset (*C* = 6.4 Torr) reduces the 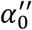 of values closest to zero PO_2_ and simplifies its physiological interpretation. Data are presented in the originally published units and the points in panels A and B are color-coded to match.

The final PO_2_, *α*_0_ curve morphologies are ones in which *α*_0_ increases continuously throughout the entire trial (Fig. 6) or peaks and then decreases at low PO_2_. These types of profiles may be generated either by calibration error or by valid physiological processes. A decrease in *α*_0_ at low PO_2_ likely indicates some physiological failure or metabolic suppression in a hypoxic environment and not calibration error. If *S* is sufficiently large relative to P_c_, the plateau in *α*_0_ described earlier may not be observed and correcting the data would produce a better estimate of *α* compared to the uncorrected value.

**Figure 6.**
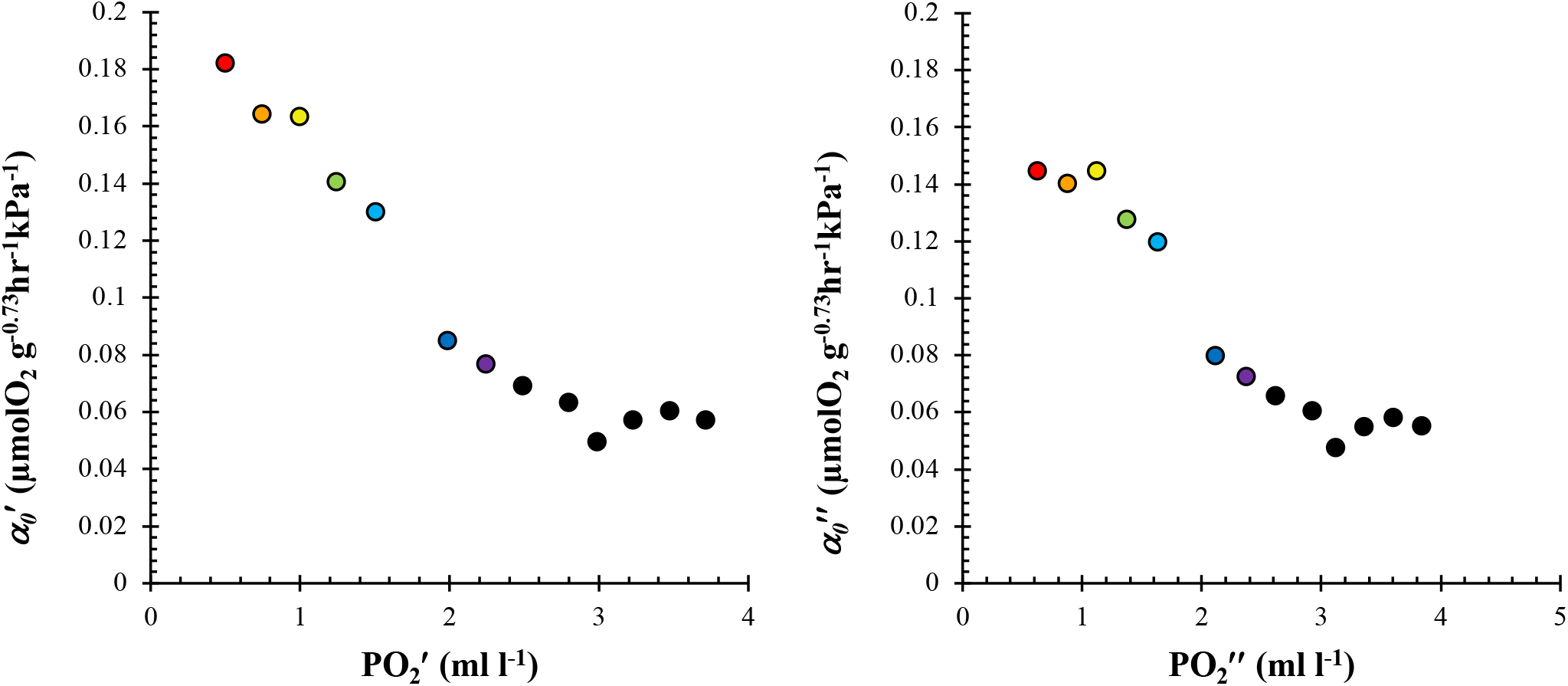
An example PO_2_, *α* curve for the goldfish *Carassius auratus* (Prosser et al., 1957) illustrating **A)** a continuous increase in *α* with falling PO_2_ and **B)** the result of correction (*C* = 0.0106 ml l^-1^). Profiles of this type are possible if *S* is sufficiently large relative to P_c_ or if the animal did not reach P_c_ during the trial. Thus, correcting profiles of this type is at the researcher’s discretion. If Pc was not reached, correcting the data will not improve estimates of α. Data in panels **A** and **B** are color-coded to match and are presented in the originally published units.

Alternatively, continuously increasing *α*_*0*_ may indicate that the animal never reached *α* during the respirometry trial. In such a case, correcting the data would be inadvisable as it would fail to generate a physiologically meaningful result 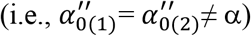. Due to this ambiguity, the decision to correct trials with profiles of this type is left to the researcher’s best judgement. If one is confident that *α* was reached, or if *α* is unrealistically high, correction may be appropriate. If a trial does not show evidence of calibration error, *C* = 0.

Many datasets will have multiple solutions to eq. 14, especially if there are negative oxygen pressures (PO_2_′ < 0). In these cases, select the smallest *C* such that PO_2_” no longer satisfies a correction criterion as described above. For example, consider a respirometry trial for which some PO_2_′ < 0. If the smallest solution to eq. 14 results in 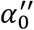 that plateau at intermediate PO_2_” and subsequently increase at the lowest-recorded PO_2_”, the next-largest solution to eq. 14 may be appropriate.

#### Nitrogen Test Experiments

Nitrogen was bubbled through seawater to mimic an animal operating at MMR and allow us to compare corrected data to a standard without calibration error. Because bubbling and stirring rates differed between trials, all *α* values were normalized per trial to reference probes without calibration error. An example curve demonstrating the effect of correction is provided in Appendix 2. Correcting data as described above substantially improved the pseudo-α calculated from probes with calibration error (Fig. 7). Because many improperly-calibrated probes recorded negative PO_2_ during trials, *α* in the uncorrected dataset is elevated and inconsistent compared to the standard. However, post-correction the mean *α* is on average only 1% higher than the reference values from properly-calibrated probes. There is no significant difference between corrected values and *α* measured using probes without calibration error, (Welch test; *t* = 1.63; df = 34; p > 0.05), though the variance is higher in corrected results (F-test; df=15, 19; *F* = 8.75; p < 0.005). Correction values are significantly correlated to known error (Spearman’s D; p < 0.001; Fig. 8), but the absolute magnitudes are different. For the range tested, correction values underestimate error by an average by 19% but provide similar estimates of alpha post-correction. This difference is not surprising, as random error in probe readings may add additional complexity to estimating the “true” correction value *C*. While any synergistic effects of random and systematic error on *α* appear to be accounted for post-correction, further investigation is warranted.

**Figure 7.**
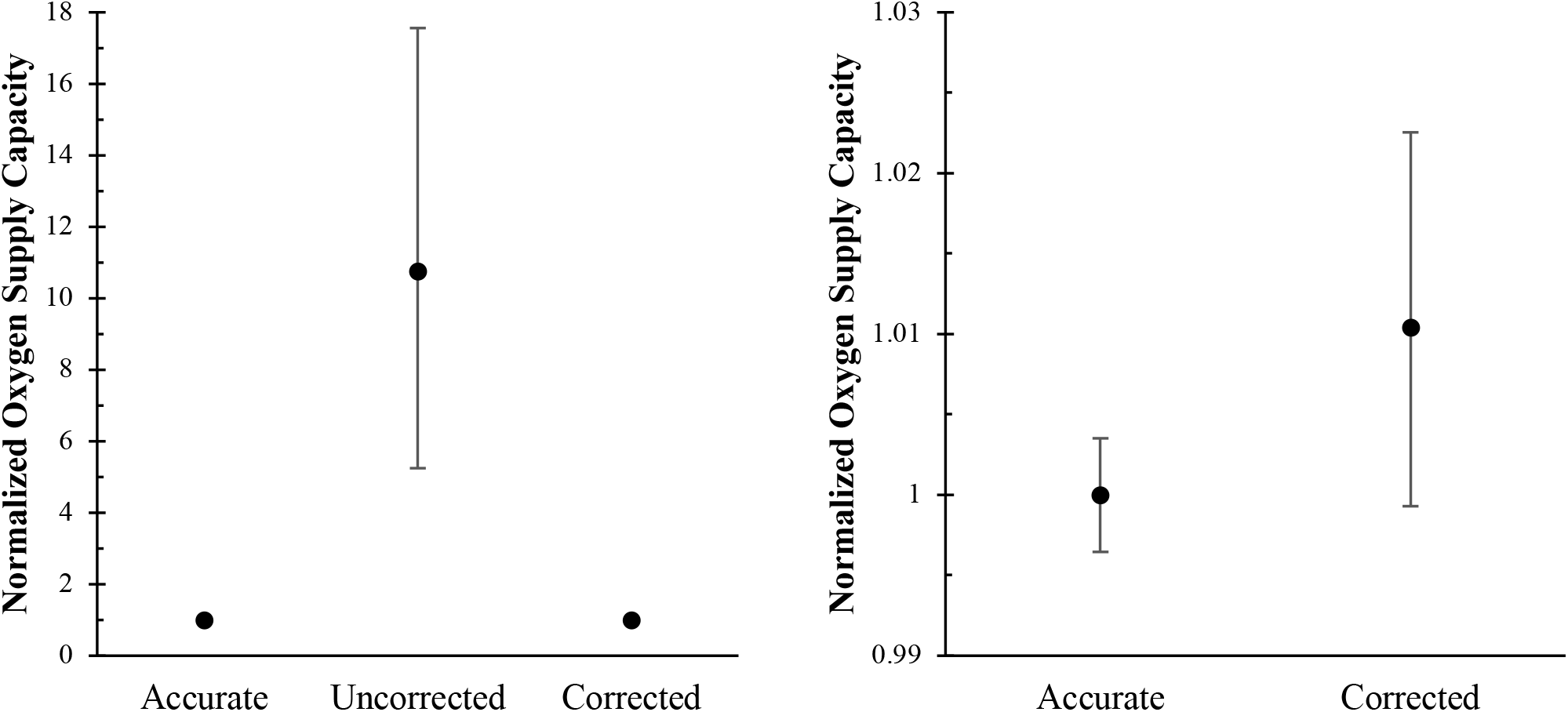
**A)** The effects of correction on nitrogen test experiments. Accurate values are the mean oxygen supply capacity of accurately-calibrated probes normalized by the mean *α* of reference probes in each trial. The low variance of this value indicates probes without error give consistent results. “Uncorrected” values are the calculated *α* of probes with calibration error normalized by the mean of reference probes. “Corrected” values are the *α* of the improperly-calibrated probes post-correction, normalized by the reference value. All values represent the mean ± 95% bootstrapped CI with n=100,000 iterations. **B)** The same data in **A**, with uncorrected data removed to improve visualization. Note the y-axis variation is only 0.04 units.

**Figure 8.**
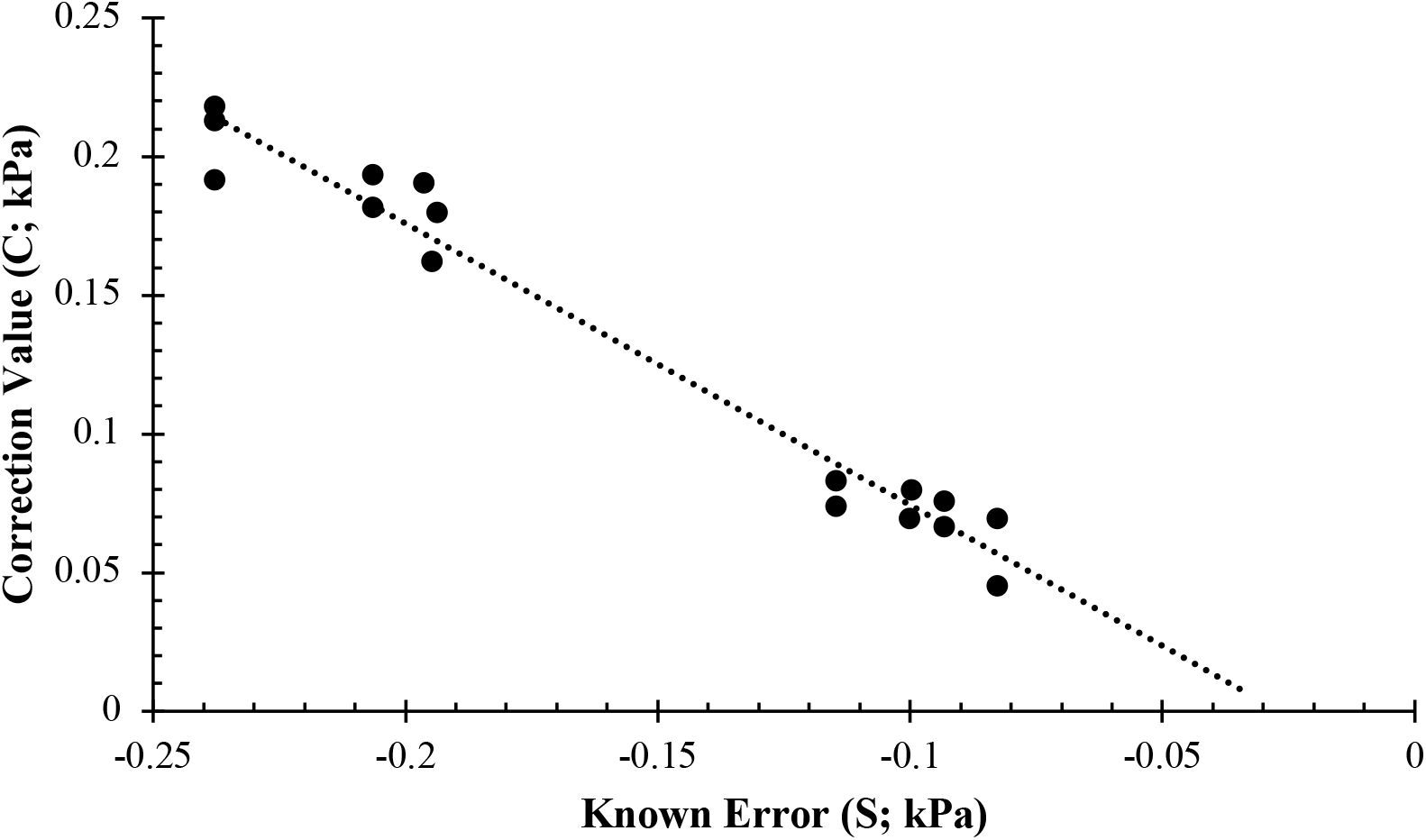
Correction value as a function of known error in nitrogen test experiments (*C* = -1.015 * *S* - 0.027; R^2^ = 0.97; *p* < 0.0001). The slope of the regression is negative because C = -S (eq. 6a).

#### Evaluation using literature datasets

To evaluate the proportion of respirometry trials where a correction for systematic error may be necessary, we conducted a survey of the literature for datasets that could be extracted for reanalysis. This resulted in 152 respirometry trials from 81 species spanning six phyla (Table 2). Of the trials surveyed, 16% (*n* = 24) met a definite correction criterion; either a plateau in *α*_*0*_ at mid-PO_2_ followed by an increase at low PO_2_ (“Plateau, increase” in Table 2) or negative PO_2_ values. Another 21% (*n* = 33) may suffer from calibration error or, alternatively, never reached *α* (or P_c_) as indicated by a continuous increase in *α*_*0*_ with falling PO_2_ (“Continuous increase” in Table 2). The remaining 63% (*n* = 95) did not meet any criteria for correction. The number of trials requiring correction is likely inflated due to inconsistent treatment of MR averaging period among experiments. For many trials, data are either sparse with very few observations reported below P_c_ (i.e., long averaging periods or very short experiments), or had abundant data calculated using relatively small averaging periods that are more susceptible to random probe error. Additionally, many manuscripts report only averaged MR values from several trials at common PO_2_ values, complicating the interpretation of C if many calibration errors are averaged together or if the reported PO_2_ values are not the average oxygen concentration of data used to calculate each MR.

**Table 2.** Calculated (*α′*) and corrected (*α′′*) oxygen supply capacity from published datasets, converted from the original published units into (μmol O_2_ g^-1^ hr^-1^ kPa^-1^). Correction values and criteria are provided for trials that may require correction. Valid criteria for correction include negative PO_2_ or a plateau in *α*_*0*_ at intermediate PO_2_ accompanied by an increase near anoxia (Plateau, increase). Trials during which *α*_*0*_ continuously increases as PO_2_ decreases (*α* incr. to zero) may require correction or may show that P_c_ was never reached.

| Species Name | Common Name | T (°C) | $\alpha'$ | Corr. Req'd | Criteria Met | C (kPa) | $\alpha''$ | Comments |
| --- | --- | --- | --- | --- | --- | --- | --- | --- |
| *Acantheephyra curtirostris* | Peaked shrimp | 5.5 | 1.547 | Yes | Plateau, increase | 0.157 | 1.279 | (Childress, 1975) |
| *Anuropus bathypelagicus* | Bathypelagic isopod | 5.5 | 1.747 | No | None | 0 | 1.747 | (Childress, 1975) |
| *Bathycalanus bradyi* | Calanoid copepod | 4 | 2.884 | Maybe | $\alpha$ incr. to zero | 0.736 | 0.906 | (Childress, 1975) |
| *Boreomysis californica* | Mysid shrimp | 5.5 | 3.583 | No | None | 0 | 3.583 | (Childress, 1975) |
| *Callinectes sapidus* | Blue crab | 17 | 0.636 | No | None | 0 | 0.636 | (Brill et al., 2015) |
| *Callinectes sapidus* | Blue crab | 23 | 0.336 | No | None | 0 | 0.336 | (Brill et al., 2015) |
| *Carcinus maenas* | European green crab | 18 | 0.164 | No | None | 0 | 0.164 | (Taylor et al., 1977)*; full seawater |
| *Carcinus maenas* | European green crab | 10 | 0.124 | No | None | 0 | 0.124 | (Taylor et al., 1977)*; full seawater |
| *Carcinus maenas* | European green crab | 18 | 0.194 | No | None | 0 | 0.194 | (Taylor et al., 1977)*; half seawater |
| *Carcinus maenas* | European green crab | 10 | 0.162 | No | None | 0 | 0.162 | (Taylor et al., 1977)*; half seawater |
| *Elenacalanus princeps* | Calanoid copepod | 5.5 | 5.499 | Maybe | $\alpha$ incr. to zero | 0.283 | 2.996 | (Childress, 1975) |
| *Euphausia pacifica* | Pacific krill | 5.5 | 0.222 | No | None | 0 | 0.222 | (Childress, 1975) |
| *Eusergestes similis* | Sergestoid decapod | 6 | 1.189 | No | None | 0 | 1.189 | (Childress, 1975) |
| *Fagegnathophausia gracilis* | Lophogastrid crustacean | 4 | 0.789 | Maybe | $\alpha$ incr. to zero | 0.415 | 0.507 | (Childress, 1975) |
| *Gaussia princeps* | Calanoid copepod | 5.5 | 1.355 | Maybe | $\alpha$ incr. to zero | 0.324 | 0.861 | (Childress, 1975) |
| *Gennadas propinquus* | Panaeoid decapod | 5.5 | 1.257 | No | None | 0 | 1.257 | (Childress, 1975) |
| *Gigantocypris agassizi* | Giant ostracod | 4 | 0.577 | No | None | 0 | 0.577 | (Childress, 1975) |
| *Gnathophausia zoea* | Lophogastrid crustacean | 5.5 | 0.787 | No | None | 0 | 0.787 | (Childress, 1975) |
| *Hymenodora frontalis* | Pacific ambereye | 5.5 | 0.630 | No | None | 0 | 0.630 | (Childress, 1975) |
| *Hyperia galba* | Big-eye amphipod | 5.5 | 1.786 | Yes | Plateau, increase | 0.185 | 1.050 | (Childress, 1975) |
| *Jasus Edwardsii* | Southern rock lobster | 13 | 0.137 | Yes | Plateau, increase | 0.856 | 0.089 | (Crear and Forteach, 2000) |
| *Jasus Edwardsii* | Southern rock lobster | 13 | 0.151 | No | None | 0 | 0.151 | (Crear and Forteach, 2000) |
| *Lithodes santolla* | Southern king crab | 12 | 0.645 | Maybe | $\alpha$ incr. to zero | 0.619 | 0.491 | (Paschke et al., 2010) |
| *Neognathophausia gigas* | Lophogastrid crustacean | 5.5 | 0.706 | No | None | 0 | 0.706 | (Childress, 1975) |
| *Neognathophausia ingens* | Giant red mysid | 5.5 | 10.73 | No | None | 0 | 10.73 | (Childress and Seibel, 1998); "Maximum Activity" |
| *Neognathophausia ingens* | Giant red mysid | 5.5 | 14.61 | Yes | Plateau, increase | 0.097 | 10.50 | (Childress and Seibel, 1998)* |
| *Neognathophausia ingens* | Giant red mysid | 5.5 | 9.141 | Maybe | $\alpha$ incr. to zero | 0.027 | 7.924 | (Childress and Seibel, 1998); non-swimming animal |
| *Notostomus, spp.* | Acantheephyrid decapod | 5.5 | 0.836 | Maybe | $\alpha$ incr. to zero | 0.986 | 0.522 | (Childress, 1975) |
| *Pasiphaea chacei* | Pasiphaeid decapod | 7.5 | 1.253 | No | None | 0 | 1.253 | (Childress, 1975) |
| *Pasiphaea emarginata* | Pasiphaeid decapod | 5.5 | 4.774 | Maybe | $\alpha$ incr. to zero | 0.595 | 0.639 | (Childress, 1975) |
| *Pasiphaea pacifica* | Pacific glass shrimp | 7.5 | 2.244 | Maybe | $\alpha$ incr. to zero | 0.072 | 1.968 | (Childress, 1975) |
| *Penaeus esculentus* | Brown tiger prawn | 25 | 0.710 | Maybe | $\alpha$ incr. to zero | 0.553 | 0.598 | (Dall, 1986); Active |
| *Penaeus esculentus* | Brown tiger prawn | 25 | 2.281 | No | None | 0 | 2.281 | (Dall, 1986); Single, 12.3g |
| *Penaeus esculentus* | Brown tiger prawn | 25 | 1.486 | No | None | 0 | 1.486 | (Dall, 1986); Single, 13.8g |
| *Penaeus esculentus* | Brown tiger prawn | 25 | 1.053 | Maybe | $\alpha$ incr. to zero | 0.787 | 0.596 | (Dall, 1986); Single, 23g |
| *Penaeus esculentus* | Brown tiger prawn | 25 | 0.789 | Maybe | $\alpha$ incr. to zero | 0.602 | 0.579 | (Dall, 1986); Inactive |
| *Phronima sedentaria* | Phronimid amphipod | 10 | 0.560 | No | None | 0 | 0.560 | (Childress, 1975) |
| *Plesionika, spp.* | Pandalid shrimp | 5.5 | 0.796 | No | None | 0 | 0.796 | (Childress, 1975) |
| *Pseudocallisoma coecum* | Scopelocherid amphipod | 5.5 | 2.112 | Maybe | $\alpha$ incr. to zero | 0.125 | 1.715 | (Childress, 1975) |
| *Sergia phorca* | Sergestoid decapod | 5.5 | 1.317 | No | None | 0 | 1.317 | (Childress, 1975) |
| *Systellapsis cristata* | Constant prawn | 5.5 | 1.259 | Maybe | $\alpha$ incr. to zero | 0.995 | 0.552 | (Childress, 1975) |
| *Ammodytes tobianus* | Sand-eel | 10 | 0.591 | Yes | Plateau, increase | 0.311 | 0.477 | (Behrens and Steffensen, 2007)* |
| *Anarhichas minor* | Spotted wolffish | 8 | 0.174 | No | None | 0 | 0.174 | (Claireaux and Chabot, 2016); 1139g |
| *Anarhichas minor* | Spotted wolffish | 8 | 0.190 | No | None | 0 | 0.190 | (Claireaux and Chabot, 2016); 2712g |
| *Aptychotrema rostrata* | Shovelnose ray | 28 | 0.342 | No | None | 0 | 0.342 | (Speers-Roesch et al., 2012)* |
| *Carassius auratus* | Goldfish | 22 | 23.62 | Yes | Plateau, increase | 0.403 | 21.91 | (Prosser et al., 1957) |
| *Carassius auratus* | Goldfish | 22 | 18.85 | Yes | Plateau, increase | 2.581 | 10.41 | (Prosser et al., 1957) |
| *Carassius auratus* | Goldfish | 22 | 20.66 | Maybe | $\alpha$ incr. to zero | 0.333 | 18.53 | (Prosser et al., 1957) |
| *Carassius auratus* | Goldfish | 22 | 13.44 | No | None | 0 | 13.44 | (Prosser et al., 1957) |
| *Chitala ornata* | Clown knifefish | 27 | 0.244 | No | None | 0 | 0.244 | (Tuong et al., 2018)* |
| *Chitala ornata* | Clown knifefish | 33 | 0.235 | No | None | 0 | 0.235 | (Tuong et al., 2018)* |
| *Crinia georgiana* | Quacking frog | 12 | 0.019 | No | None | 0 | 0.019 | (Marshall et al., 2013)<sup>o</sup> |
| *Crinia georgiana* | Quacking frog | 12 | 0.063 | Maybe | $\alpha$ incr. to zero | 0.537 | 0.020 | (Mueller and Seymour, 2011)<sup>+</sup> |
| *Cryptobranchus alleganiensis* | Hellbender salamander | 20 | 0.120 | No | None | 0 | 0.120 | (Yeager and Ultsch, 1989); 100g |
| *Cyclopterus lumpus* | Lumpfish | 10 | 0.415 | No | None | 0 | 0.415 | (Ern et al., 2016)* |
| *Cyclopterus lumpus* | Lumpfish | 16 | 0.543 | No | None | 0 | 0.543 | (Ern et al., 2016)* |
| *Cymatogaster aggregata* | Shiner perch | 14 | 1.262 | No | None | 0 | 1.262 | (Snyder et al., 2016) |
| *Cymatogaster aggregata* | Shiner perch | 14 | 1.469 | No | None | 0 | 1.469 | (Snyder et al., 2016) |
| *Danio rerio* | Zebrafish | 28 | 19.04 | Yes | Plateau, increase | 0.234 | 15.60<br>0 | (Mandic et al., 2020)*; 10 days post fertilization |
| *Danio rerio* | Zebrafish | 28 | 12.22 | No | None | 0 | 12.21<br>7 | (Mandic et al., 2020)*; 15 days post fertilization |
| *Danio rerio* | Zebrafish | 28 | 12.61 | No | None | 0 | 12.60<br>6 | (Mandic et al., 2020)*; 4 days post fertilization |
| *Danio rerio* | Zebrafish | 28 | 13.91 | No | None | 0 | 13.91<br>3 | (Mandic et al., 2020)*; 7 days post fertilization |
| *Danio rerio* | Zebrafish | 28 | 4.206 | No | None | 0 | 4.206 | (Mandic et al., 2020)*; adult |
| *Fundulus grandis* | Gulf killifish | 27 | 1.638 | No | None | 0 | 1.638 | (Reemeyer and Rees, 2019); closed respirometry |
| *Fundulus grandis* | Gulf killifish | 27 | 1.728 | No | None | 0 | 1.728 | (Reemeyer and Rees, 2019); closed respirometry |
| *Fundulus grandis* | Gulf killifish | 27 | 1.732 | No | None | 0 | 1.732 | (Reemeyer and Rees, 2019); intermittent respirometry |
| *Gadus morhua* | Atlantic cod | 5 | 0.301 | No | None | 0 | 0.301 | (Schurmann and Steffensen, 1997) |
| *Hemiscyllium ocellatum* | Epauvette shark | 28 | 0.388 | No | None | 0 | 0.388 | (Speers-Roesch et al., 2012)* |
| *Hypseleotris, spp.* | Carp gudgeon | 25 | 0.966 | No | None | 0 | 0.966 | (Stoffels, 2015) |
| *Lates calcarifer* | Barramundi | 26 | 1.010 | Yes | Plateau, increase | 0.889 | 0.642 | (Collins et al., 2013); Broome sub-population |
| *Lates calcarifer* | Barramundi | 36 | 1.875 | Yes | Plateau, increase | 0.789 | 1.162 | (Collins et al., 2013); Broome sub-population |
| *Lates calcarifer* | Barramundi | 26 | 0.566 | No | None | 0 | 0.566 | (Collins et al., 2013);<br>Darwin sub-population |
| *Lates calcarifer* | Barramundi | 36 | 1.882 | No | None | 0 | 1.882 | (Collins et al., 2013);<br>Darwin sub-population |
| *Lates calcarifer* | Barramundi | 26 | 0.822 | No | None | 0 | 0.822 | (Collins et al., 2013);<br>Gladstone sub-population |
| *Lates calcarifer* | Barramundi | 36 | 2.172 | Maybe | $\alpha$ incr.<br>to zero | 0.291 | 1.793 | (Collins et al., 2013);<br>Gladstone sub-population |
| *Lumpenus lampraeformis* | Snake blenny | 15 | 0.325 | No | None | 0 | 0.325 | (Pelster et al., 1988) |
| *Lumpenus lampraeformis* | Snake blenny | 15 | 0.413 | No | None | 0 | 0.413 | (Pelster et al., 1988) |
| *Mayaheros urophthalmus* | Nile chichlid | 23 | 1.276 | No | None | 0 | 1.276 | (Burggren et al.,<br>2019a)* |
| *Mayaheros urophthalmus* | Nile chichlid | 28 | 2.225 | Maybe | $\alpha$ incr.<br>to zero | 1.523 | 0.737 | (Burggren et al.,<br>2019a)* |
| *Mayaheros urophthalmus* | Nile chichlid | 33 | 1.929 | Maybe | $\alpha$ incr.<br>to zero | 0.135 | 1.637 | (Burggren et al.,<br>2019a)* |
| *Melanocetus johnsonii* | Humpback<br>anglerfish | 28 | 0.089 | No | None | 0 | 0.089 | (Cowles and<br>Childress, 1995)* |
| *Melanostigma pammelas* | Midwater<br>eelpout | 5.5 | 0.750 | No | None | 0 | 0.750 | (Childress, 1975) |
| *Melanotaenia fluviatilis* | Australian<br>rainbowfish | 25 | 1.569 | No | None | 0 | 1.569 | (Stoffels, 2015) |
| *Mogurnda adspersa* | Purple-<br>spotted<br>gudgeon | 25 | 0.879 | No | None | 0 | 0.879 | (Stoffels, 2015) |
| *Nectoliparis pelagicus* | Tadpole<br>snailfish | 5.5 | 1.116 | No | None | 0 | 1.116 | (Childress, 1975) |
| *Necturus maculosus* | Common<br>mudpuppy | 12 | 0.184 | Yes | Plateau,<br>increase | 1.402 | 0.117 | (Marshall et al.,<br>2013) |
| *Necturus maculosus* | Common<br>mudpuppy | 20 | 0.299 | Yes | Plateau,<br>increase | 0.300 | 0.203 | (Yeager and Ultsch,<br>1989); 126g |
| *Necturus maculosus* | Common<br>mudpuppy | 20 | 0.200 | Yes | Plateau,<br>increase | 1.751 | 0.111 | (Yeager and Ultsch,<br>1989); 129g |
| *Opsanus tau* | Oyster<br>toadfish | 22 | 0.381 | Yes | Plateau,<br>increase | 1.088 | 0.240 | (Ultsch et al., 1981);<br>310g |
| *Opsanus tau* | Oyster<br>toadfish | 22 | 0.327 | No | None | 0 | 0.327 | (Ultsch et al., 1981);<br>314g |
| *Opsanus tau* | Oyster<br>toadfish | 22 | 0.332 | No | None | 0 | 0.332 | (Ultsch et al., 1981);<br>349g |
| *Opsanus tau* | Oyster<br>toadfish | 22 | 0.291 | No | None | 0 | 0.291 | (Ultsch et al., 1981);<br>364g |
| *Opsanus tau* | Oyster<br>toadfish | 22 | 0.297 | Yes | Plateau,<br>increase | 0.052 | 0.280 | (Ultsch et al., 1981);<br>395g |
| *Opsanus tau* | Oyster<br>toadfish | 22 | 0.382 | Yes | Plateau,<br>increase | 0.729 | 0.252 | (Ultsch et al., 1981);<br>437g |
| *Oreochromis niloticus* | Nile tilapia | 30 | 0.375 | No | None | 0 | 0.375 | (Burggren et al.,<br>2019b)* |
| *Paralichthys dentatus* | Summer flounder | 22 | 0.144 | Maybe | $\alpha$ incr. to zero | 0.416 | 0.131 | (Capossela et al., 2012)*; Acclimated to 22°C |
| *Paralichthys dentatus* | Summer flounder | 30 | 0.163 | No | None | 0 | 0.163 | (Capossela et al., 2012)*; Acute exposure to 30°C |
| *Perca fluviatilis* | European perch | 20 | 0.655 | No | none | 0 | 0.655 | (Thuy et al., 2010)*; *, after sham feeding |
| *Perca fluviatilis* | European perch | 20 | 0.807 | Maybe | $\alpha$ incr. to zero | 1.058 | 0.551 | (Thuy et al., 2010)*; *, before sham feeding |
| *Perca fluviatilis* | European perch | 20 | 0.500 | No | None | 0 | 0.500 | (Thuy et al., 2010)*; *, fasting |
| *Perca fluviatilis* | European perch | 20 | 0.813 | Yes | Plateau, increase | 0.114 | 0.749 | (Thuy et al., 2010)*; force feeding |
| *Pseudophryne bibronii* | Brown toadlet | 12 | 0.027 | No | None | 0 | 0.027 | (Mueller and Seymour, 2011)<sup>+</sup> |
| *Reinhardtius hippoglossoides* | Greenland halibut | 5 | 0.383 | No | None | 0 | 0.383 | (Claireaux and Chabot, 2016); 72g; |
| *Reinhardtius hippoglossoides* | Greenland halibut | 5 | 0.408 | No | None | 0 | 0.408 | (Claireaux and Chabot, 2016); 89g |
| *Reinhardtius hippoglossoides* | Greenland halibut | 5 | 0.249 | No | None | 0 | 0.249 | (Claireaux and Chabot, 2016); 2.391 kg |
| *Savelinus fontinalis* | Brook trout | 12 | 0.906 | No | None | 0 | 0.906 | (Claireaux and Chabot, 2016); 10.5g; D. Chabot, unpub. Data |
| *Sciaenops ocellatus* | Red drum | 24 | 1.392 | No | None | 0 | 1.392 | (Ern et al., 2016)* |
| *Sciaenops ocellatus* | Red drum | 30 | 1.608 | No | None | 0 | 1.608 | (Ern et al., 2016)* |
| *Sciaenops ocellatus* | Red drum | 26 | 1.603 | No | None | 0 | 1.603 | (Negrete and Esbaugh, 2019)*; closed respirometry |
| *Sciaenops ocellatus* | Red drum | 26 | 1.504 | No | None | 0 | 1.504 | (Negrete and Esbaugh, 2019)*; intermittent respirometry |
| *Siren lacertna* | Greater siren | 20 | 0.056 | Yes | Plateau, increase | 1.683 | 0.037 | (Yeager and Ultsch, 1989); 388g |
| *Hemipholis elongata* | Brittle star | 20 | 1.062 | Maybe | $\alpha$ incr. to zero | 1.328 | 0.351 | (Christensen and Colacino, 2000)* |
| *Hemipholis elongata* | Brittle star | 20 | 1.049 | Yes | Plateau, increase | 1.025 | 0.424 | (Christensen and Colacino, 2000)*; hyperpnea |
| *Abrialiopsis felis* | Enoploteuthid squid | 5 | 0.823 | No | None | 0 | 0.823 | (Hunt and Seibel, 2000) |
| *Abrialiopsis pacificus* | Enoploteuthid squid | 5 | 0.797 | No | None | 0 | 0.797 | Hunt & Seibel (2000) |
| *Argopecten purpuratus* | Purple scallop | 25 | 0.377 | Yes | Plateau, increase | 0.247 | 0.311 | (Aguirre-Velarde et al., 2016) |
| *Argopecten purpuratus* | Purple scallop | 16 | 0.245 | Yes | Plateau, increase | 1.073 | 0.120 | (Aguirre-Velarde et al., 2016) |
| *Bathyteuthis abyssicola* | Deepsea squid | 5 | 0.574 | No | None | 0 | 0.574 | (Seibel et al., 1997) |
| *Doryteuthis (Amerigo) pealeii* | Longfin squid | 15 | 0.832 | No | None | 0 | 0.832 | (Birk et al., 2018) |
| *Doryteuthis (Amerigo) pealeii* | Longfin squid | 15 | 1.317 | No | None | 0 | 1.317 | (Birk et al., 2018) |
| *Doryteuthis (Amerigo) pealeii* | Longfin squid | 15 | 1.309 | No | None | 0 | 1.309 | (Birk et al., 2018) |
| *Doryteuthis (Amerigo) pealeii* | Longfin squid | 15 | 1.087 | No | None | 0 | 1.087 | (Birk et al., 2018) |
| *Dosidicus gigas* | Jumbo squid | 23-27 | 19.07 | Maybe | $\alpha$ incr. to zero | 0.342 | 5.099 | (Birk et al., 2018) |
| *Dosidicus gigas* | Jumbo squid | 23-27 | 6.911 | Maybe | $\alpha$ incr. to zero | 0.802 | 4.080 | (Birk et al., 2018) |
| *Dosidicus gigas* | Jumbo squid | 23-27 | 7.308 | Yes | Plateau, increase | 0.894 | 4.482 | (Birk et al., 2018) |
| *Dosidicus gigas* | Jumbo squid | 23-27 | 6.265 | Maybe | $\alpha$ incr. to zero | 1.123 | 3.159 | (Birk et al., 2018) |
| *Gibberulus gibbosus* | Humpbacked conch | 28 | 1.691 | Maybe | $\alpha$ incr. to zero | 0.033 | 1.334 | Lefevre et al., 2015)*; ambient CO2 |
| *Gibberulus gibbosus* | Humpbacked conch | 33 | 4.057 | Maybe | $\alpha$ incr. to zero | 0.194 | 1.253 | (Lefevre et al., 2015)*; ambient CO2 |
| *Gonatus onyx* | Clawed armhook squid | 5 | 2.050 | No | None | 0 | 2.050 | (Hunt and Seibel, 2000); juvenile |
| *Gonatus onyx* | Clawed armhook squid | 5 | 1.078 | No | None | 0 | 1.078 | (Hunt and Seibel, 2000); sub-adult |
| *Histioteuthis heteropsis* | Strawberry squid | 5 | 0.176 | No | None | 0 | 0.176 | (Seibel et al., 1997) |
| *Japetella diaphana* | Pelagic octopod | ? | 0.592 | Maybe | $\alpha$ incr. to zero | 0.364 | 0.198 | (Birk et al., 2019) |
| *Japetella diaphana* | Pelagic octopod | 5 | 0.044 | No | None | 0 | 0.044 | (Seibel et al., 1997) |
| *Japetella heathi* | Pelagic octopod | 5 | 0.139 | Yes | Plateau, increase | 0.24 | 0.074 | (Seibel et al., 1997) |
| *Laternula elliptica* | Antarctic clam | 0 | 0.726 | Maybe | $\alpha$ incr. to zero | 3.818 | 0.219 | (Peck et al., 2002) |
| *Laternula elliptica* | Antarctic clam | 0 | 0.308 | No | None | 0 | 0.308 | (Peck et al., 2002) |
| *Laternula elliptica* | Antarctic clam | 0 | 0.229 | No | None | 0 | 0.229 | (Peck et al., 2002) |
| *Laternula elliptica* | Antarctic clam | 0 | 0.314 | No | None | 0 | 0.314 | (Peck et al., 2002) |
| *Laternula elliptica* | Antarctic clam | 0 | 0.475 | No | None | 0 | 0.475 | (Peck et al., 2002) |
| *Nautilus pompilius* | Chambered nautilus | 25 | 0.316 | Yes | Plateau, increase | 1.368 | 0.270 | (Redmond et al., 1978) |
| *Nautilus pompilius* | Chambered nautilus | 11 | 0.093 | No | None | 0 | 0.093 | (Staples et al., 2000) |
| *Nautilus pompilius* | Chambered nautilus | 16 | 0.218 | No | None | 0 | 0.218 | (Staples et al., 2000) |
| *Nautilus pompilius* | Chambered nautilus | 21 | 0.110 | No | None | 0 | 0.110 | (Staples et al., 2000) |
| *Octopus bimaculoides* | Two-spot octopus | 10 | 0.235 | No | None | 0 | 0.235 | (Seibel and Childress, 2000) |
| *Octopus bimaculoides* | Two-spot octopus | 10 | 0.198 | No | None | 0 | 0.198 | (Seibel and Childress, 2000) |
| *Octopus bimaculoides* | Two-spot octopus | 10 | 0.336 | Maybe | $\alpha$ incr. to zero | 0.048 | 0.285 | (Seibel and Childress, 2000) |
| *Octopus californicus* | California octopus | 6 | 0.181 | No | None | 0 | 0.181 | (Seibel and Childress, 2000) |
| *Octopus californicus* | California octopus | 6 | 0.239 | No | None | 0 | 0.239 | (Seibel and Childress, 2000) |
| *Panopea zelandica* | New Zealand geoduck | 15 | 16.75 | Maybe | $\alpha$ incr. to zero | 0.091 | 4.382 | (Le et al., 2016) |
| *Vampyroteuthis infernalis* | Vampire squid | 5 | 0.016 | No | None | 0 | 0.016 | (Seibel et al., 1997) |
| *Sipunculus nudus* | Peanut worm | 15 | 0.235 | Maybe | $\alpha$ incr. to zero | 0.475 | 0.086 | (Pörtner et al., 1985)*; Large animals |
| *Sipunculus nudus* | Peanut worm | 15 | 0.283 | Maybe | $\alpha$ incr. to zero | 0.361 | 0.100 | (Pörtner et al., 1985)*; small animals |

\* Data originally presented as mean ± error. Values extracted are the mean.
+ No mass units provided in manuscript; assumed to be g<sup>-1</sup>

The effect of the correction method on replicate measures of *α* within a species was assessed using respirometry data from the OMZ copepod *Megacalanus* (Wishner et al., 2018) measured at 5ºC and 10ºC. Due to these animals’ low P_c_, all trials approached anoxia and therefore probe calibration error would likely have an outsized effect on oxygen supply capacity. Of the 16 trials evaluated, 9 recorded negative PO_2_ and a total of 12 showed evidence of calibration error. Correcting data reduced both the mean and variance of both datasets, and post-correction there is a significant difference in oxygen supply capacity at 5ºC and 10ºC (ANOVA; *df*_*t*_ = 15, *F* = 103.7, *p* < 0.001; Fig. 9). Clearly, correcting for calibration error has implications for data interpretation and has conceptual value by explaining extreme outliers in some datasets like this one.

**Figure 9.**
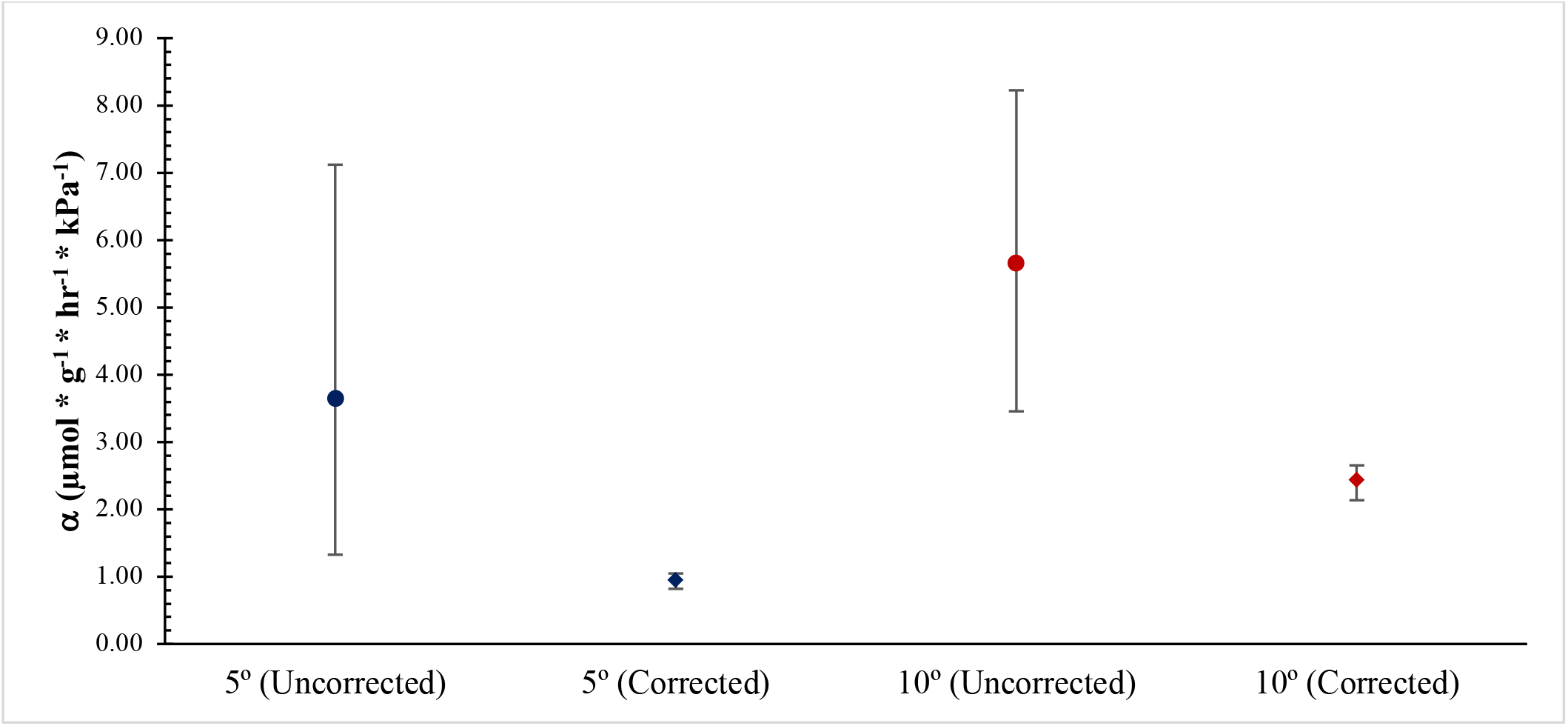
Oxygen supply capacity (*α*) for the OMZ copepod *Megacalanus* from Wishner et al. (2018) at 5ºC (n = 9) and 10ºC (n = 7) pre (uncorrected) and post-correction. Values represent the mean ± 95% bootstrapped CI with n = 100,000 iterations. Correction values ranged from 0.005 to 0.19 kPa (mean = 0.10 kPa).

## Conclusions

The method of Seibel et al., (2021b), used to determine the oxygen supply capacity (*α*), facilitates the identification and correction of systematic error in PO_2_ measurement. When appropriate, adding a small PO_2_ correction value to each observation that equalizes the two highest measures of oxygen provision (*α*_0_) counteracts the effects of this error and provides a better estimate of *α* and other metabolic traits.

There are two definitive scenarios in which this correction method should be employed: 1) if negative PO_2_ values are recorded, or 2) if there is a plateau in measured oxygen supply (*α*_0_′) at intermediate PO_2_ followed by a dramatic increase at the lowest recorded oxygen levels. Alternative relationships between *α*_0_′ and PO_2_may represent valid physiological processes and not error. If no calibration error is suspected, no correction is merited and C = 0. Our literature survey found that approximately 16% of published datasets meet our proposed criteria for correction. Validation experiments with nitrogen showed that, post-correction, improperly-calibrated oxygen probes provide comparable estimates of *α* to properly-calibrated reference probes. Correcting a dataset from the OMZ copepod *Megacalanus* where some trials showed evidence of calibration error reduced both the mean and variance of α, in part by reducing extreme, high outliers. The need for data correction does not indicate shoddy, improper, or haphazard work. It is entirely possible, and indeed probable, for small amounts of systematic error to appear in datasets despite a researcher’s diligent efforts and use of best practices. While precise calibration and high-resolution equipment can reduce the magnitude of such error, it is impossible to eliminate completely and should thus be accounted for when analyzing respirometry data.

## Summary

1. This correction method successfully mitigates the effect of systematic PO_2_ error caused by inaccurate probe calibration on some measured metabolic traits.
2. Small amounts of calibration error can affect measures of oxygen supply capacity and P_c_ and must be accounted for, especially in species that live in low-oxygen environments where the effect of this error is magnified.
3. When multiple, valid potential correction values are identified, we recommend using the smallest one such that, after correction, the trial no longer meets a correction criterion. Valid correction values are typically positive and reflect erroneous calibration at anoxia.
4. For the data analyzed here, the correction method reduced variability among replicate observations of oxygen supply capacity (and consequently P_c_ calculated as P_c_ = MR/α).
5. Based on a literature survey, a correction criterion is met in approximately one of six published respirometry curves.
6. For trials that do not show evidence of calibration error, the correction value C = 0.

## Acknowledgements

This project was supported by the Garrels Memorial Fellowship and William and Elsie Knight Endowed Fellowship for Marine Science to AWT, and NSF grant OCE-2127538 and NOAA grant UWSC11082 to BAS.

## Appendix 1: Reported oxygen probe characteristics

**Table A1:**
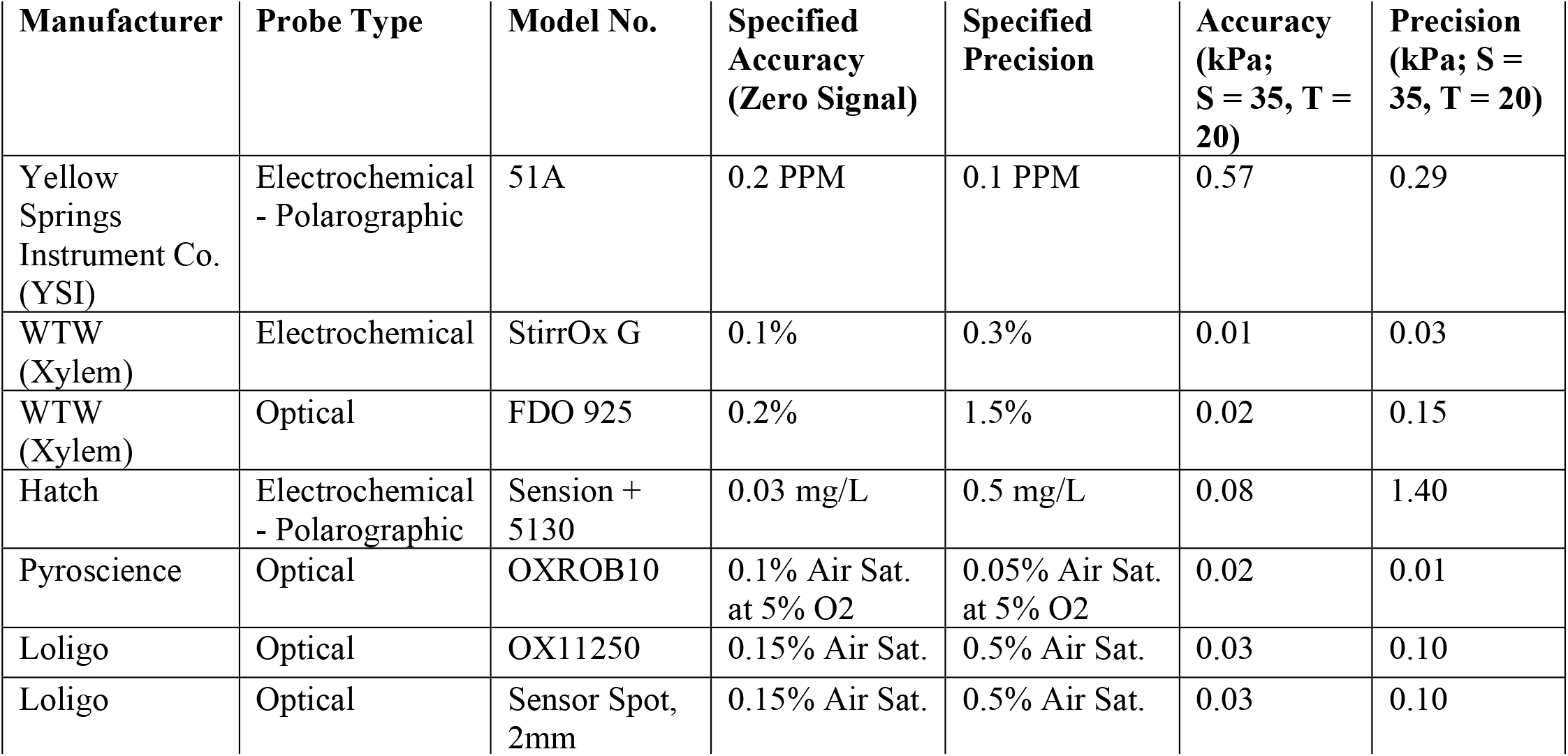
Accuracy at zero oxygen and precision (repeat measurements) for several different types of oxygen probes as reported in the manufacturer specification sheets. Values were standardized to kPa in seawater at 20ºC and a salinity (S) of 35.

| Manufacturer | Probe Type | Model No. | Specified Accuracy (Zero Signal) | Specified Precision | Accuracy (kPa; S = 35, T = 20) | Precision (kPa; S = 35, T = 20) |
| --- | --- | --- | --- | --- | --- | --- |
| Yellow Springs Instrument Co. (YSI) | Electrochemical - Polarographic | 51A | 0.2 PPM | 0.1 PPM | 0.57 | 0.29 |
| WTW (Xylem) | Electrochemical | StirrOx G | 0.1% | 0.3% | 0.01 | 0.03 |
| WTW (Xylem) | Optical | FDO 925 | 0.2% | 1.5% | 0.02 | 0.15 |
| Hatch | Electrochemical - Polarographic | Sension + 5130 | 0.03 mg/L | 0.5 mg/L | 0.08 | 1.40 |
| Pyroscience | Optical | OXROB10 | 0.1% Air Sat. at 5% O<sub>2</sub> | 0.05% Air Sat. at 5% O<sub>2</sub> | 0.02 | 0.01 |
| Loligo | Optical | OX11250 | 0.15% Air Sat. | 0.5% Air Sat. | 0.03 | 0.10 |
| Loligo | Optical | Sensor Spot, 2mm | 0.15% Air Sat. | 0.5% Air Sat. | 0.03 | 0.10 |

## Appendix 2: Example MMR dataset with error

**Figure A2.1.**
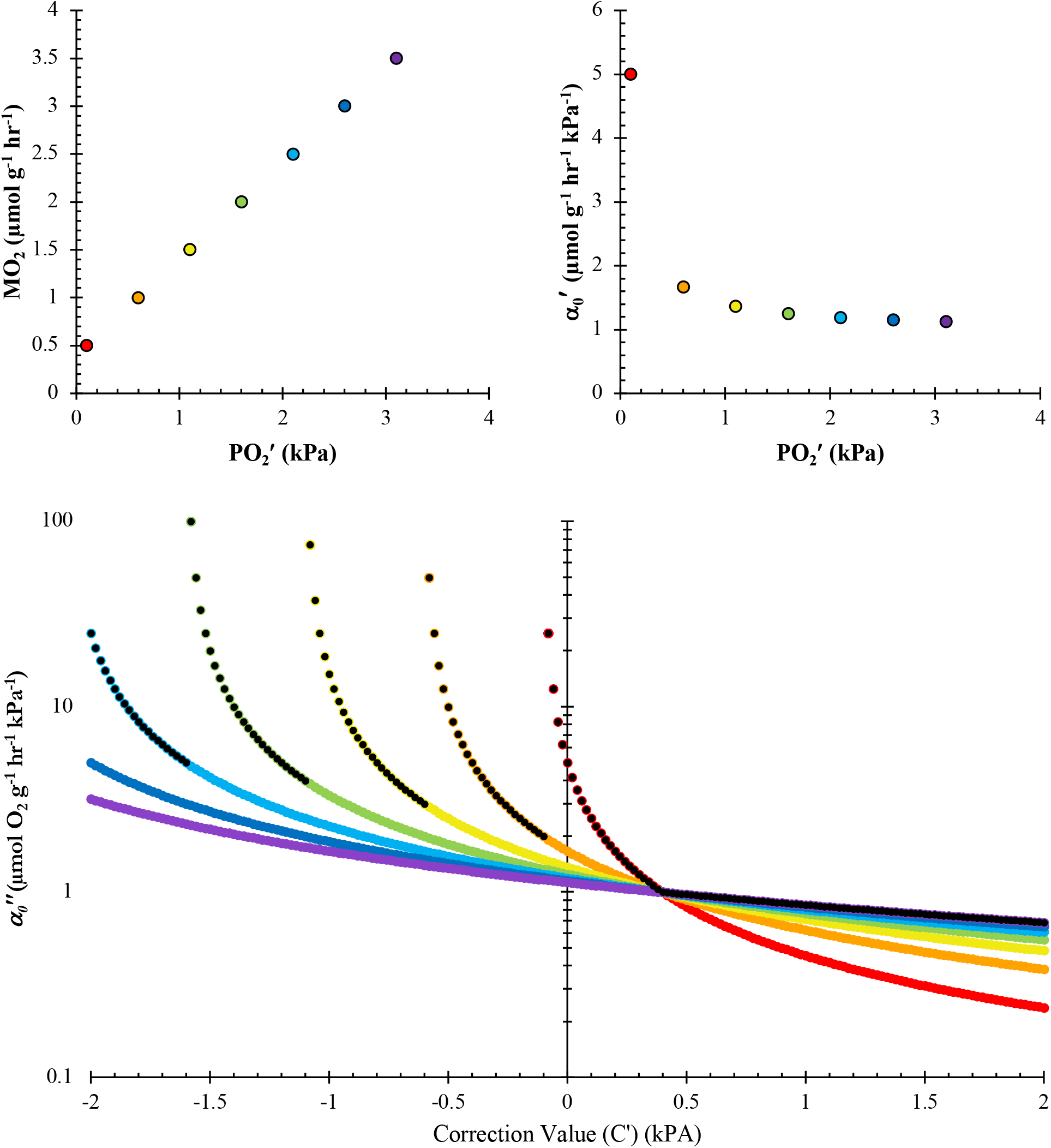
**A)** Example maximum metabolic rate (MMR) dataset {(MO_2(*n*)_, PO_2_*′* _(*n*)_), …, (MO_2(*N*)_, PO_2_*′* _(*N*)_)} with error S = -0.4 kPa. The selected *α* for this dataset = 1.0. **B)** 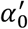 as a function of 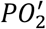. **C)** 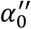 as a function of Δ*C′* at intervals Δ*C′* = 0.02 kPa for all points in the dataset in panel **A** and are color-coded to match. The superimposed black dots identify the points used to define the piecewise function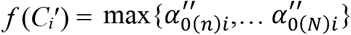. A corrected supply measurement 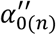 becomes negative when *C′* > PO_2(n)_ and is not displayed on a log-scale. Note that, in this dataset where *S =* -0.4, the only solution to 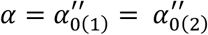 occurs at *C′* = *C* = -*S* = 0.4. The trial satisfies a correction criterion, as 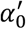 increases dramatically at the lowest-observed PO_2_.

Note as the sum of the error term *S* and PO_2_ approaches zero (i.e., PO_2_ + *S* → 0), the denominator in (eq 5) approaches zero. Evaluating limits illustrates how 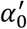 diverges as *S* approaches -PO_2_:

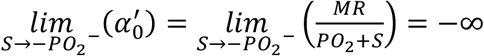

and

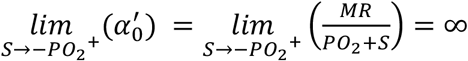

We are most concerned with right-sided limit where 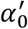 diverges to infinity, as it can dramatically raise the estimate of *α* and thereby over-estimate the predicted maximum achievable MR at any environmental PO_2_ and decrease the estimation of P_c_ for a given MR.

**Figure A2.2.**
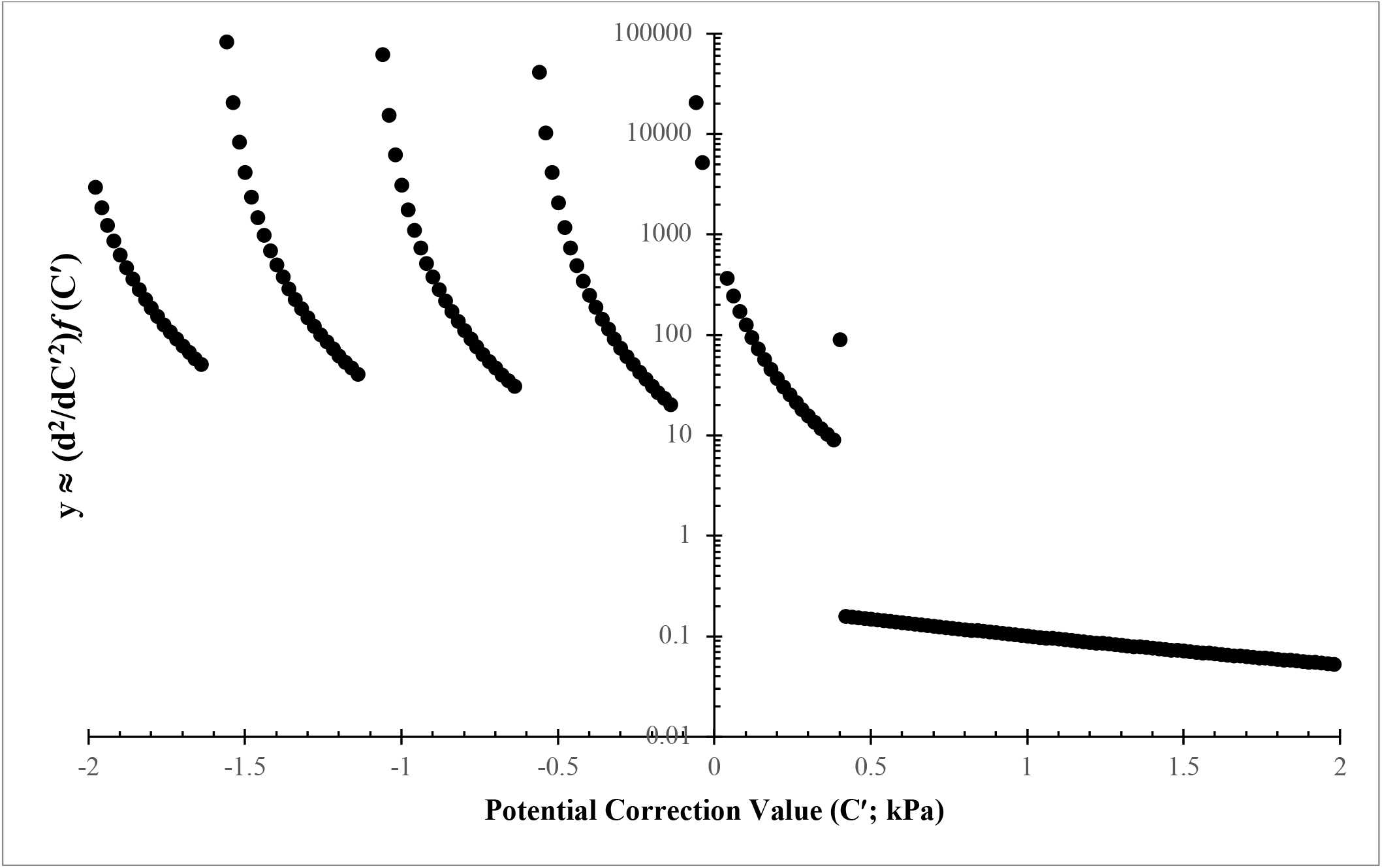
Second-order difference quotient of with respect to for the MMR dataset presented in Fig. A1. Discontinuities where potential *C′*= *C* = −*S* are identified by a large difference (∼doubling) between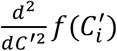 and 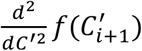 or 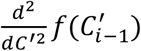, and where 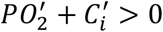 for all 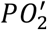. For this dataset where = 0., the only appropriate discontinuity occurs at *C′* = 0.4. Discontinuities at negative would result in some 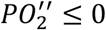 and are therefore invalid.

## Appendix 3: Example reference (072122_3_1) and probe with error (072122_3_3) from nitrogen trials

**Figure A2.3:**
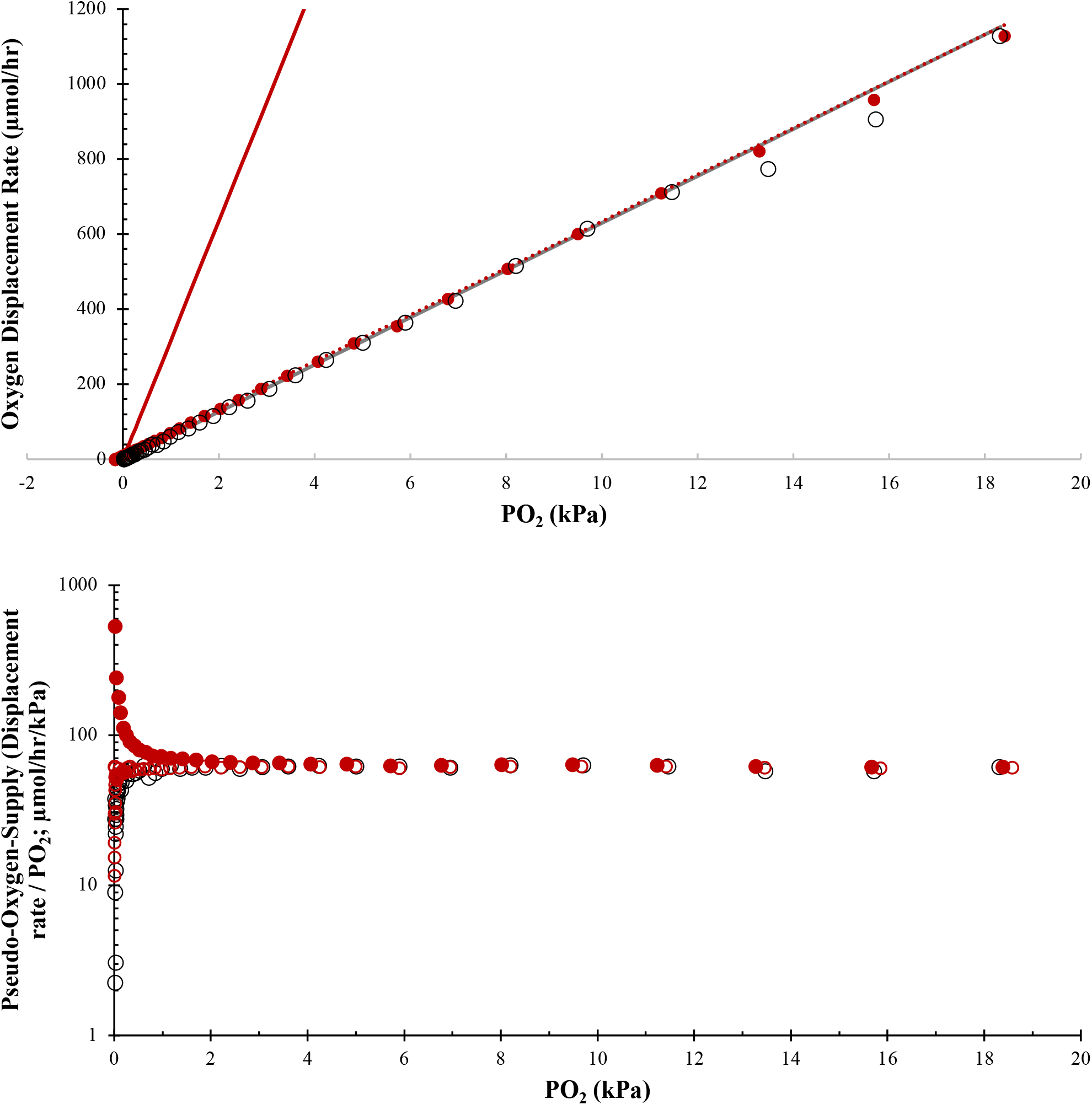
Effect of calibration error and the correction method on nitrogen-derived data. **A)** Oxygen displacement rate (μmol/hr; analogous to MO_2_) as a function of PO_2_. Red circles are data from a probe with a known calibration error S = -0.0194 ± 0.0011 kPa (mean ± SD) and the reference probe (open black circles, S = 0.0017 ± 0.0011 kPa; mean ± SD). The α-line, used to calculate the maximum achievable aerobic metabolic rate at any environmental PO_2_, is shown for the uncorrected data (solid red line; *α* = 317 μmol/hr/kPa), corrected data (dashed red line; C = 0.18 kPa; *α* = 62.3 μmol/hr/kPa), and the reference data (gray line; *α* = 62.9 μmol/hr/kPa). **B)** Oxygen displacement rate divided by PO_2_ (pseudo-*α*_0_) as a function of PO_2_ for the reference probe (black open circles), uncorrected data (closed red circles), and corrected data (open red circles; C = 0.18 kPa). Note the log-scale y-axis and how the uncorrected observations increase dramatically near anoxia.

## Appendix 4: What if calibration error is a function of PO_2_?

Many probes have two-point calibrations where the sensor value at air saturation (100% PO_2_) and anoxia (0% PO_2_) are both recorded. Therefore, at intermediate PO_2_ values, it is reasonable to suspect that error in the 0% calibration (S), which causes divergence in *α*_0_ near anoxia, is weighted according to some function of PO_2_. Here we refer to this weighting function as and the effective error at some intermediate oxygen concentration is equal to the weighting function, evaluated at that PO_2_, multiplied by the error at anoxia (*S*).

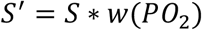

For example, if a perfect 100% saturation calibration is performed, a simple linear weighting function could be defined where

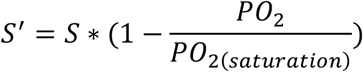

According to this function, systematic error is nil at 100% saturation (i.e., zero-point calibration error does not affect observations at air saturation) and effective error increases linearly to S as PO_2_ decreases to anoxia. If we assume a non-constant weighting function, equation 3 is invalid and MR becomes a function of error, PO_2_, and the weighting function and is consequently it is not possible to identify as described in the manuscript. If this is the case, the correction method can only be used in an iterative process that is significantly more complicated than the method, as described. To estimate with a nonconstant weighting function, A potential correction value, 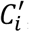, one of many from a set of values hypothesized *a priori* that may satisfy *S* +*C* = 0, is assessed. The effective potential correction value 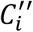 for each PO_2_ is created when 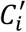 is weighted by the assumed weighting function. MR is then calculated from modified data:

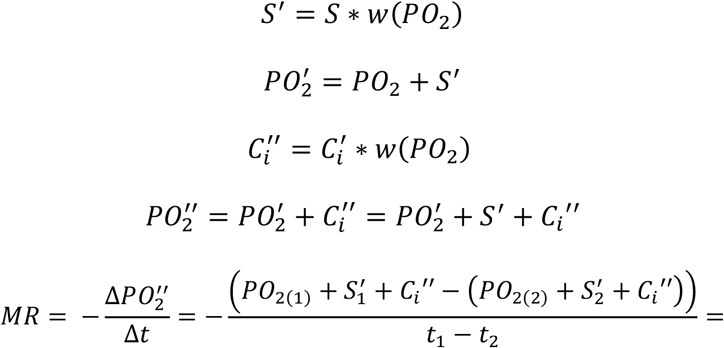

To find valid correction values, effective potential correction values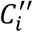, must be added iteratively to raw data (PO_2_ as a function of *t*) and the “corrected” MR for each potential correction value is calculated from the “corrected” PO_2_ data. Next, 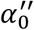 for each point in the 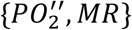 dataset is calculated by dividing “corrected” MR by 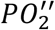 and the highest 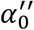 is identified. A function analogous to eq. 11 can then be written to describe 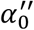 as a function of *C′*. Valid potential correction values (i.e., where *C′*= *C* = −*S*) could similarly be identified using discontinuities in the second-order difference quotient, analogous to as described in eq. 13a. The value of these discontinuities could then be iteratively by narrowing the range of tested until sufficient precision, decided by the researcher, is achieved.

Calculating MR assuming a non-constant weighting function increases the difficulty of estimating C and provides no practical benefit compared to assuming S is constant for all PO_2_. First and foremost, an alternative weighting function must be assumed that captures S as a function of PO_2_. While this could potentially be measured empirically by systematically creating an array of incorrect calibrations (as described in our N_2_ experiments) and testing the probes with error against a reference probe across a PO_2_ gradient, it is not guaranteed to be constant in all conditions, across time as the sensor ages, or across all oxygen probes. Additionally, because S is usually small relative to the total change in PO_2_ during an experiment and because bins used to calculate MR are usually much less than the length of a trial, the differential effect on MR between a constant and nonconstant weighting function is insignificant in the context of respirometry experiments. For example, within an averaging bin, if ΔPO_2_ from the raw trace is 1 kPa and Δt is 1 hr, ignoring any normalizations for chamber volume and units, MR calculated with constant S is 1 kPa/hr. If we assume the linear weighting function as described above, we must adjust PO_2_ first before calculating MR. In this case where S = -0.1kPa, PO_2(2)_ is 0.0048 kPa lower than it “should” be and, after adjustment, the real MR is corrected to 0.9952 kPa/hr; 0.48% or 1 part in 210 different than the value calculated using a constant S. This variability is much less than any reported variability in SMR of individuals of the same species we could find in the literature and is much less than the typical variation in MR within an experiment. Moreover, most respirometry trials attempt to measure *α* (or P_c_) at SMR and therefore the *α* used to calculate *α* have similar PO_2_. Because observations have similar PO_2_ (i.e., observations close to saturation and close to anoxia are not typically compared), any analytical errors introduced by assuming an invariant *S* are minimized.

With these considerations in mind, we find the myriad mathematical simplicities and practical benefits provided by assuming a constant weighting function outweigh any theoretical drawbacks. Therefore, we recommend using the constant weighting function of S implied in this manuscript.

