## Supplementary material for "A method for correcting systematic error in PO_2_ measurement to improve measures of oxygen supply capacity": correct_alpha.R

```
correct_alpha <- function(po2, mo2,cRange = 2, discBreak = 2, results = 1,
                          view_c = NULL, rawplot = 0, diffplot = 0){
  
# Function to calculate PO2 correction values for respirometry data
# ------
  
# Arguments:
# po2       <- a vector of PO2 values
# mo2       <- a vector of MO2 values. Must be the same length and corresponding
#              to PO2
# cRange    <- input C range (~10% pO2 at saturation) [default = 2; kPa]
# discBreak <- log(DiscBreak) is the cutoff value for identifying discontinuities
#              in the second derivative [default = 2]. 
# results   <- optionally display valid C and the corresponding a0" in a table
#              (no table is produced if C=0. [default = 1])
# view_c    <- view_c = n; optionally display a0" vs PO2" for the n smallest values 
#              of C identified within cRange (no plots provided if view_c = n =0;
#              [default n = NULL produces a plot for every C in cRange])
# rawplot   <- optionally display a plot of uncorrected alpha v po2
#              (plot generated if rawplot !=0 [default = 0])
# diffplot  <- optionally display a plot of the abs(log(second order difference
#              )) vs cPrime (plot generated if diffplot != 0 [default = 0])

# Outputs:
# a0_Prime    <- maximum a0' of the uncorrected dataset
# C           <- list of valid C > 0
# a0_dblPrime <- maximum a0" for each valid C

# Author: Alexander W. Timpe, 2020
#--------

# Load ggplot2
require(ggplot2)
library(ggplot2)
  
# Force po2 and mo2 as vectors
po2_untrimmed <- as.vector(po2)
mo2_untrimmed <- as.vector(mo2)

po2 <- po2_untrimmed[!is.na(po2_untrimmed)&!is.na(mo2_untrimmed)]
mo2 <- mo2_untrimmed[!is.na(po2_untrimmed)&!is.na(mo2_untrimmed)]

# Verify that po2 and mo2 have the same number of observations
if(length(po2)!=length(mo2)){
  stop("po2 and mo2 vectors must be the same length")
}

# Determine the maximum alpha in the uncorrected dataset
a0prime     <- mo2/po2
raw_a       <- max(a0prime,na.rm = TRUE)

# Create a matrix of potential correction values, cPrime
cPrime      <- -c(seq(from = -abs(cRange),to = abs(cRange), by = cRange/100))

# Preallocate a results matrix for corrected PO2 values
cPrimeMxa   <- matrix(0,nrow=length(mo2),ncol=length(cPrime))

# Apply each Cprime(i) to each po2
for (i in 1:ncol(cPrimeMxa)){
  cPrimeMxa[,i]      <- (po2+cPrime[i])
  }

# Determine the a0prime for each PO2 for each Cprime(i)
cPrimeMxb   <- mo2/cPrimeMxa

# Preallocaate a results matrix for max(CPrime(i))
ma02Prime   <- matrix(0,ncol=ncol(cPrimeMxb))

# Determine the maximum a0prime for each Cprime(i)
ma02Prime   <- apply(cPrimeMxb,2,max)

# Preallocate a results matrix for the second-order difference quotient
SecOrdDiff  <- matrix(0,nrow=length(as.vector(ma02Prime))-2)
SecOrdDiff  <- cbind(cPrime[2:200],SecOrdDiff)

# Determine the second-order difference quotient for each max(a0prime)
for (i in 1:nrow(SecOrdDiff)){
  SecOrdDiff[i,2]    <- (ma02Prime[i]-2*ma02Prime[i+1]+ma02Prime[
                                i+2])/((abs(cPrime[i]-cPrime[i+1]))^2)
  }

# Take the base 10 logarithm of each second-order difference quotient
PosSODiff   <- abs(SecOrdDiff[,2])
logPSOD     <- log10(PosSODiff)

# Preallocate a results matrix for identifying discontinuities in the
# second derivative of a0prime with respect to C
LRRL                 <- matrix(0,nrow=nrow(SecOrdDiff),ncol=2)
colnames(LRRL)       <- c("LR","RL")

# Determine valid discontinuities
for (i in 1:(nrow(LRRL)-1)){
  LRRL[i+1,1] <- logPSOD[i]/logPSOD[i+1]
  LRRL[i,2]   <- logPSOD[i+1]/logPSOD[i]
  }
VDiscIdx1   <- abs(LRRL[,"RL"]-1) > log10(discBreak)
VDiscIdx2   <- abs(LRRL[,"LR"]-1) > log10(discBreak)
VDiscIdx3   <- cPrime[2:200]+min(po2) > 0
VDiscIdx4   <- (VDiscIdx1+VDiscIdx2)>=1 & VDiscIdx3>0

# Verify that valid discontinuities were identified
if(sum(VDiscIdx4)==0){
  # Generate the list of outputs and assign values
  resList             <- list()
  resList$a0_Prime    <- NA
  resList$C           <- NA
  resList$a0_dblPrime <- NA
  paste("No valid discontinuities identified. Modify cRange to continue")
  return(resList)
  }else{
  
  # Set aside the C' of valid discontinuities
  VDiscPO2    <- cPrime[2:200]
  VDiscr      <- sort(VDiscPO2[VDiscIdx4])
  VDisc       <- VDiscr[1:(length(VDiscr)-1)]
  VDisc       <- c(0,VDisc)
  
  # Save the untrimmed PO2 data for the raw plot later
  rdat        <- as.data.frame(cbind(po2,a0prime))
  
  # Remove negative MO2 values and the PO2 values associated with them
  mo2idx      <- mo2>=0
  pdat_mx     <- cbind(po2,mo2)
  pdat_mxP    <- pdat_mx[mo2idx,]
  mo2         <- pdat_mxP[,"mo2"]
  po2         <- pdat_mxP[,"po2"]
  
  # Create a matrix for potential correction values and their stable equilibria
  Cmx         <- matrix(0,nrow=length(VDisc),ncol=2)
  Cmx[,1]     <- VDisc  
  
  # Determine stable discontinuities
  for(j in 1:length(VDisc)){
    testC           <- VDisc[j]
    testPo2         <- po2+testC
    testamax        <- max(mo2/testPo2)
    a0dblprm        <- mo2/testPo2
    a0dblprmSort    <- sort(a0dblprm,TRUE)
    a0dpIdx1        <- a0dblprm == a0dblprmSort[1]
    a0dpIdx2        <- a0dblprm == a0dblprmSort[2]
    po2dpr_1        <- po2[a0dpIdx1]
    po2dpr_2        <- po2[a0dpIdx2]
    po2dpr_1        <- po2dpr_1[1]
    po2dpr_2        <- po2dpr_2[1]
    c_mo2_1         <- mo2[a0dpIdx1]
    c_mo2_2         <- mo2[a0dpIdx2]
    c_mo2_1         <- c_mo2_1[1]
    c_mo2_2         <- c_mo2_2[1]
    C               <- ((c_mo2_1*po2dpr_2)-(c_mo2_2*po2dpr_1))/(c_mo2_2-c_mo2_1)
    if(is.nan(C)){
      Cmx[j,2]      <- NaN
    }else if(C+min(po2)<0){
      Cmx[j,2]      <- NaN
    }else Cmx[j,2]  <- C
  }
  
  # Extract valid discontinuities
  CmxIdx              <- !is.nan(Cmx[,2])
  Valid_Disc          <- Cmx[CmxIdx,1]
  
  # Verify that valid discontinuities were identified
   if(length(Valid_Disc)==0){
     resList             <- list()
     resList$a0_Prime    <- NA
     resList$C           <- NA
     resList$a0_dblPrime <- NA
     paste("No valid discontinuities identified. Modify cRange to continue")
     return(resList)
     }else{
    
    # Determine stable correction values
    Cmx2               <- matrix(0,nrow=length(Valid_Disc),ncol=6)
    Cmx2[,1]           <- Valid_Disc
    
    for(i in 1:5){
      for(j in 1:nrow(Cmx2)){
        testC           <- Cmx2[j,i]
        testPo2         <- po2+testC
        testamax        <- max(mo2/testPo2)
        a0dblprm        <- mo2/testPo2
        a0dblprmSort    <- sort(a0dblprm,TRUE)
        a0dpIdx1        <- a0dblprm == a0dblprmSort[1]
        a0dpIdx2        <- a0dblprm == a0dblprmSort[2]
        a0pr_1          <- min(a0dblprm[a0dpIdx1])
        a0pr_2          <- max(a0dblprm[a0dpIdx2])
        po2dpr_1        <- min(po2[a0dpIdx1])
        po2dpr_2        <- max(po2[a0dpIdx2])
        c_mo2_1         <- min(mo2[a0dpIdx1])
        c_mo2_2         <- max(mo2[a0dpIdx2])
        C               <- ((c_mo2_1*po2dpr_2)-(c_mo2_2*po2dpr_1))/(c_mo2_2-c_mo2_1)
        Cmx2[j,i+1]     <- C
      }
    }
    
    # Extract the unique, stable correction values and formulate the results matrix 
    UniqueDisc      <- unique(Cmx2[,6])
    ResMx           <- matrix(0,ncol=2,nrow=length(UniqueDisc))
    colnames(ResMx) <- c('C','a0"')
    for(i in 1:nrow(ResMx)){
      StableC       <- UniqueDisc[i]
      StablePo2     <- po2+StableC
      StableA       <- max(mo2/StablePo2)
      ResMx[i,1]    <- StableC
      ResMx[i,2]    <- StableA
    }
    
    # Create a vector of non-negative, stable correction values. 
    CIdx       <- ResMx[,1]>=0
    C_final    <- ResMx[CIdx,1]
    
    # Create a vector of max(a0") corresponding to each stable, positive correction 
    # value
    alpha <- ResMx[CIdx,2]
    
    if(length(C_final)==0){
      C_final <- NA
    } else(if(max(C_final)<=0.25*cRange){
      print("cRange large relative to C. Recommend narrower cRange")
    })
    
    # Optionally print the results matrix in the console
    if(results!=0){
      print(ResMx)
    }
    
    # Optionally view the a0" for each C in the results matrix
    if(is.null(view_c)){
      numPlots    <- nrow(ResMx)
      for(i in 1:numPlots){
        testPO2   <- po2 + ResMx[i,1]
        a0dblpm   <- mo2/testPO2
        testmx    <- as.data.frame(cbind(testPO2,a0dblpm))
        rcnm      <- variable.names(testmx)
        test_plot <- ggplot(testmx, aes(x = testmx[,1], y = testmx[,2])) +
          geom_point(color = "black", 
                     shape = 18) + 
          labs(title   = paste('a0" vs. po2" ; C =',round(ResMx[i,1],digits=4)),
               x      = paste('PO2"; C = ',round(ResMx[i,1],digits=4)),
               y      = 'a0"') 
        print(test_plot)
      }
    }
    if(!is.null(view_c)){
    if(view_c>0){
      numPlots    <- min(view_c,nrow(ResMx))
      if(view_c>nrow(ResMx)){
        print("view_c > number of C in cRange. Maxmimum number of plots generated")
      }
      for(i in 1:numPlots){
        testPO2   <- po2 + ResMx[i,1]
        a0dblpm   <- mo2/testPO2
        testmx    <- as.data.frame(cbind(testPO2,a0dblpm))
        rcnm      <- variable.names(testmx)
        test_plot <- ggplot(testmx, aes(x = testmx[,1], y = testmx[,2])) +
          geom_point(color = "black", 
                     shape = 18) + 
          labs(title   = paste('a0" vs. po2" ; C =',round(ResMx[i,1],digits=4)),
               x      = paste('PO2"; C = ',round(ResMx[i,1],digits=4)),
               y      = 'a0"') 
        print(test_plot)
      }
    }
    }
      
    # Optionally produce a plot of a0' vs PO2'
    if (rawplot != 0) {
      raw_plot  <- ggplot(rdat, aes(x = rdat[,1], y = rdat[,2])) +
        geom_point(color = "black", 
                   shape = 18) + 
        labs(title  = "a0' vs. po2'",
             x      = "po2'",
             y      = "a0'") 
      print(raw_plot)
    }
    
    # Optionally produce a plot of f"(C') vs C
    if (diffplot != 0) {
      xdat      <- as.data.frame(cbind(cPrime[2:200],log10(PosSODiff)))
      cnm       <- variable.names(xdat)
      diff_plot <- ggplot(xdat, aes(x = xdat[,1], y = xdat[,2])) +
        geom_point(color = "black", 
                   shape = 18) + 
        labs(title  = "f''(C') vs. C'",
             x    = paste("C'; cRange =",cRange),
             y    = "log10(f''(C')") 
      print(diff_plot)
    }
     }
  }


# Generate the list of outputs and assign values
resList             <- list()
resList$a0_Prime    <- raw_a
resList$C           <- C_final
resList$a0_dblPrime <- alpha
return(resList)

# End Program
}
```
